# Cellular senescence preserves viral genome maintenance

**DOI:** 10.1101/2022.07.12.499783

**Authors:** Alexander M Pham, Luz E Ortiz, Aron E Lukacher, Hyun Jin Kwun

## Abstract

Senescent cells accumulate in the host during the aging process and are associated with age-related pathogenesis, including cancer. Although persistent senescence seems to contribute to many aspects of cellular pathways and homeostasis, the role of senescence in virus-induced human cancer is not well understood. Merkel cell carcinoma (MCC) is an aggressive skin cancer induced by a life-long human infection of Merkel cell polyomavirus (MCPyV). Here, we show that MCPyV large T (LT) antigen expression in human skin fibroblasts causes a novel nucleolar stress response followed by p21-dependent senescence and senescence-associated secretory phenotypes (SASPs) which are required for MCPyV genome maintenance. The senolytic, navitoclax, treatment resulted in decreased senescence and MCPyV genome levels, suggesting a potential therapeutic for MCC prevention. Our results uncover the mechanism of a host stress response regulating human polyomavirus genome maintenance in viral persistency, which may lead to the targeted intervention for MCC.

## Introduction

Senescence is a stable cell cycle arrest thought to be an antiproliferative cellular defense mechanism against cancer^1^ and viral replication^2^. This process can be induced through a variety of cellular stress responses, such as the DNA damage response (DDR), the nucleolar stress response (NSR), and replicative stresses^1,3^, leading to p53 activation and subsequent upregulation of p21. p21 can induce sustained G1 cell cycle arrest and senescence, thereby preventing cell cycle progression and proliferation^1^. Senescent cells have an altered phenotype compared to their non-senescent counterparts including the induction of senescence-associated secretory phenotype (SASP) genes, upregulation of anti-apoptotic genes^4^, and downregulation of various antiviral signaling pathways^5,6^.

Several studies have shown that viral infections can induce cellular senescence as part of the antiviral response^2,7^. Vesicular stomatitis virus titers were reduced when permissive cells were driven into senescence by bleomycin treatment^2^. Infection of fibroblast cells with measles virus induced cell fusion and cellular senescence, indicating the potential of senescence to limit viral replication^8^. On the other hand, there is growing evidence of viruses using senescent cells to their advantage. Influenza virus and varicella zoster virus (VZV) replication efficiency was higher in senescent cells compared to nonsenescent cells^6^. Senescent cells have also been shown to upregulate the entry receptor for dengue virus and severe acute respiratory syndrome coronavirus 2 (SARS-CoV2), allowing for enhanced viral entry and infection^9,10^. Because senescent cells are more resistant to apoptosis and have reduced antiviral signaling, this cellular environment could be conducive for persistent viral infections.

These senescent phenotypes could be exploited by MCPyV, a double stranded DNA tumor virus that causes persistent infection. MCPyV is a ubiquitous pathogen that can cause Merkel cell carcinoma (MCC), a rare but aggressive skin cancer mainly affecting immunocompromised patients^11^. The MCPyV LT antigen is required for viral genome replication^12^ and restriction of cell proliferation^13,14^. This protein is unique from LT antigens of other human polyomaviruses in that it contains a disordered MCPyV Unique Region (MUR) domain^15,16^. The MUR domain aids in establishing viral latency through ubiquitination by host E3 ubiquitin ligases^15,17–19^ that destabilizes MCPyV LT and results in inhibition of MCPyV DNA replication. Here, we investigate the mechanism of MCPyV LT-induced senescence and the novel role of a host cellular stress response during a human polyomavirus infection, which may explain a critical interplay between the host and a viral pathogen leading to cancer.

## Results

### MCPyV LT mediates cellular senescence in human fibroblast cells through its unique domain

During our study of MCPyV LT-mediated cell growth regulation^15^, we observed significant structural alterations in normal human fibroblasts BJ-hTERT expressing MCPyV LT, including enlarged nuclei and flattened morphology; features associated with cellular senescence. Because enlarged nuclei in MCC cases correlate with MCPyV positivity^20^, we explored the possibility that senescence, mediated by MCPyV LT, was an integral part of the MCPyV life cycle. To determine if MCPyV LT expression alone induces cellular senescence, we created lentiviruses encoding either MCPyV LT or SV40 LT fused with 2A self-cleaving peptide (P2A) and enhanced green fluorescent protein (eGFP) to transduce BJ-hTERT cells; an eGFP-P2A empty vector lentivirus served as a negative control (Figure 1a). Because the MUR domain is unique to MCPyV and regulates cell proliferation^15^, we asked if this domain had a role in regulating cell morphological changes and therefore, senescence. To understand the effect of the MUR domain on MCPyV LT-induced morphological changes, a MCPyV LT mutant with a deletion of the MUR domain (MCPyV LT dMUR)^15^ and a SV40 LT with the insertion of the MCPyV LT MUR domain (SV40LT+MUR)^15^ were also examined. Only the MCPyV LT-expressing cells strongly stained positive for senescence-associated beta-galactosidase (SA-β-gal), the most common senescent marker, at 14 days post-transduction. 14 days post-transduction was chosen as we observed the most morphological changes at this timepoint. SA-β-gal staining was determined by both brightfield microscopic (Figure 1b) and flow cytometric analyses (Figure 1c). Approximately 50% of MCPyV LT-expressing cells were senescent, whereas only 5% or less of cells were senescent for the remaining LT constructs or empty vector. SA-β-gal positive cells expressing MCPyV LT had a significant increase in nuclear area, approximately twice the size (> 500 μm^2^) compared to the empty vector and other LT constructs (225∼250 μm^2^) (Figure 1d). Minimal staining or morphological changes were observed in cells expressing the empty vector, SV40 LT, and SV40LT+MUR. MCPyV LTdMUR expression greatly reduced SA-β-gal staining and nuclear size, indicating the potential for the MUR domain to promote senescence only in the context of MCPyV LT. Notably, insertion of the MUR domain into SV40 LT did not promote senescence, suggesting that the MUR domain does not directly stimulate senescence and other factors are likely involved in inducing this phenotype. Analysis of mRNAs for a variety of senescence-associated secretory phenotype (SASPs), where transcripts for ANKRD1, CSF2, CXCL1, CXCL2, EDN1, IL-6, and IL-8 were markedly upregulated by MCPyV LT expression and only modestly upregulated in MCPyV LTdMUR-expressing cells compared to the eGFP control (Figure 1e), concluding that MCPyV LT induces senescence through the MUR domain.

**Figure 1:**
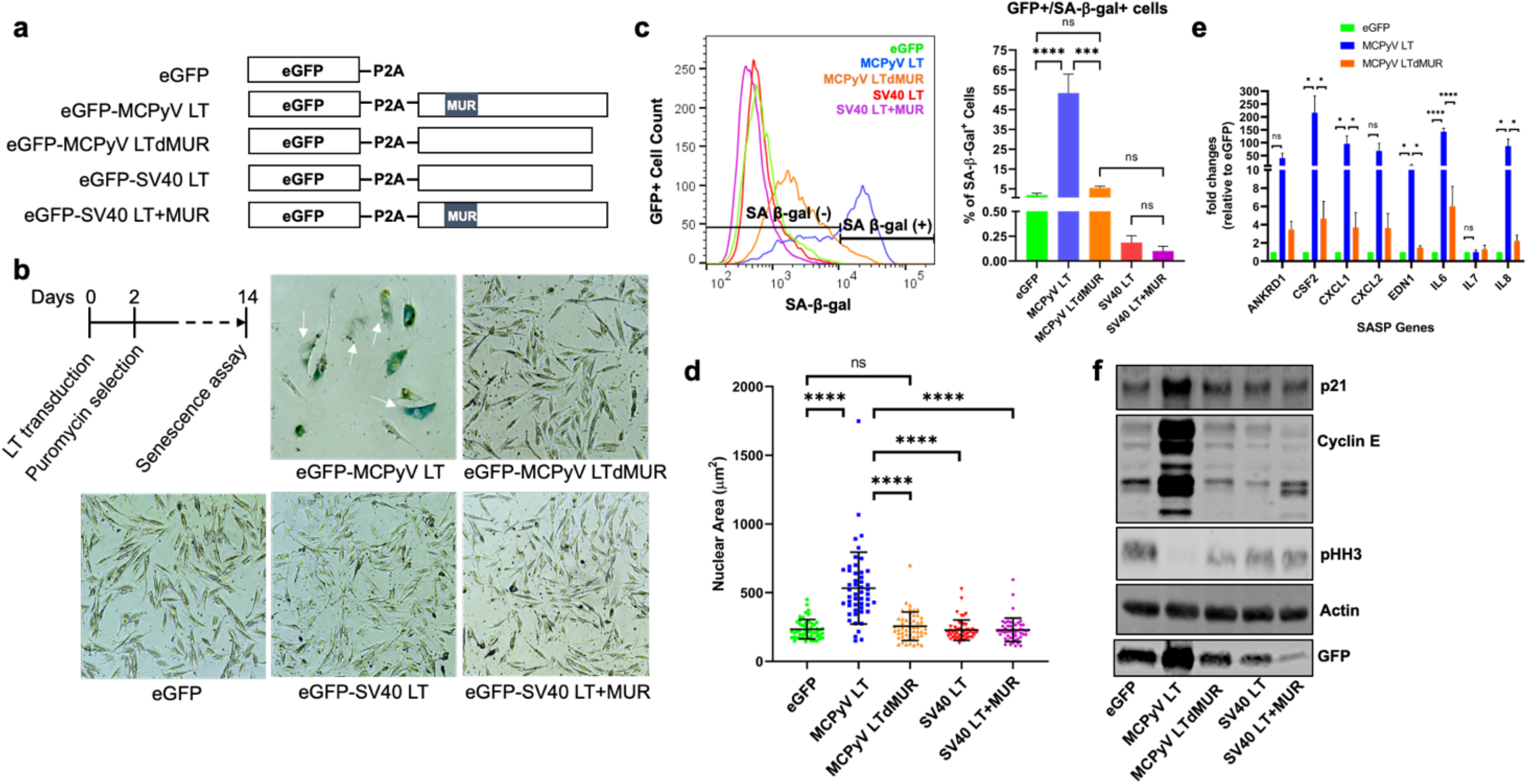
MCPyV LT induces cellular senescence. (**a**) Lentiviral constructs for LT expression. (**b**) Senescence assay. BJ-hTERT cells were transduced with either eGFP empty vector or LT constructs and were selected with puromycin at two days post transfection. A colorimetric senescence-associated β-galactosidase assay (SA-β-gal) was conducted 14 days post transduction. Cells with blue staining were positive for SA-β-gal, indicated by white arrows. Images are representative of 3 experiments. (**c**) Fluorescent intensity of SA-β-gal activity (SA-β-gal^+^) was measured in GFP^+^ (LT expressing) cells by flow cytometry. MCPyV LT expression significantly increased the number of senescent cells. One-way ANOVA test was used to determine statistical significance (*** p < 0.001, **** p < 0.0001, ns = not significant). Standard error bars represent standard error value, n = 3. (**d**) Nuclear area (µm^2^) of empty vector or LT transduced BJ-hTERT cells. MCPyV LT promotes senescent phenotypes. One-way ANOVA test (**** p < 0.0001, ns = not significant). Standard error bars represent mean value with standard deviation, n = 50 cells per construct. (e) SASP activation by MCPyV LT. Quantitative reverse transcription polymerase chain reaction (RT-qPCR) analysis of SASP cytokines. Abbreviations – ANKRD1: Ankyrin repeat domain 1, CSF2: Colony-stimulating factor 2, CXCL: chemokine (C-X-C motif) ligand, EDN1: Endothelin 1, IL: Interleukin. SASPs mRNA levels was substantially upregulated by LT expression except IL-7. One-way ANOVA test was used to determine statistical significance (**** p < 0.0001, * p < 0.05, ns = not significant). Standard error bars represent standard error value, n = 3. (f) MCPyV LT activates p21. Growth arrest genes were upregulated in MCPyV LT-expressing cells. Immunoblot of cell cycle-related genes in BJ-hTERT cells transduced with empty vector or LT constructs. Actin was used as a loading control. GFP was used to measure LT expression.

Senescence is commonly regulated by p53/p21 activation and characterized by prolonged G1 cell cycle arrest^1^. Immunoblot analysis revealed that only MCPyV LT led to p21 activation (Figure 1f). MCPyV LT-positive cells also showed substantial upregulation of cyclin E, a G1 phase marker^21–24^, indicating G1 cell cycle arrest in a subset of the cell population. In addition, histone H3 phosphorylation at Ser10 (pHH3), a mitotic marker^25^, is weakly expressed in G2 phase^26^. The MCPyV LT-positive cells displayed H3 phosphorylation at low levels, signifying cell growth arrest^27,28^. In contrast, the MCPyV LTdMUR and SV40 LT+MUR mutants showed no significant p21 upregulation nor Cyclin E or pHH3 downregulation compared to eGFP^+^ control cells, implying that growth arrest was specific to wild type MCPyV LT. Immunofluorescent analysis confirmed that enlarged MCPyV LT senescent cells exhibited high p21 expression (Figure S1a) and lacked pHH3 expression (Figure S1b). These results suggest that only MCPyV LT could activate p21, specifically through the MUR domain to induce growth arrest and cellular senescence.

### MCPyV LT-mediated senescence is NSR-dependent

Senescent growth arrest is often triggered by a persistent DNA damage response^4^. A previous study has shown that MCPyV LT can induce double stranded breaks and activate the host cell’s DDR through p53 activation^13^. To determine whether MCPyV LT induces DDR-dependent senescence, we looked for the presence of gamma-H2AX (γ-H2AX), a biomarker for DNA double-strand breaks^29^. A subset of MCPyV LT-expressing senescent cells with enlarged nuclei (>500 μm^2^) were positive for γ-H2AX staining, although some senescent cells were also negative for γ-H2AX staining despite morphologic changes (Figure 2a). Moreover, MCPyV LT-expressing cells with a normal nuclear size (∼200 μm^2^) were still able to induce a DDR response. Although eGFP, MCPyV LTdMUR, SV40 LT, and SV40 LT+MUR-expressing cells were significantly lower for SA-β-gal staining (Figure 1b, 1c), 20-30% of cells were positive for γ-H2AX foci formation regardless of the stimulus (Figure S2). These results indicate that wild-type MCPyV LT likely mediates cellular senescence through a DDR-independent manner.

**Figure 2:**
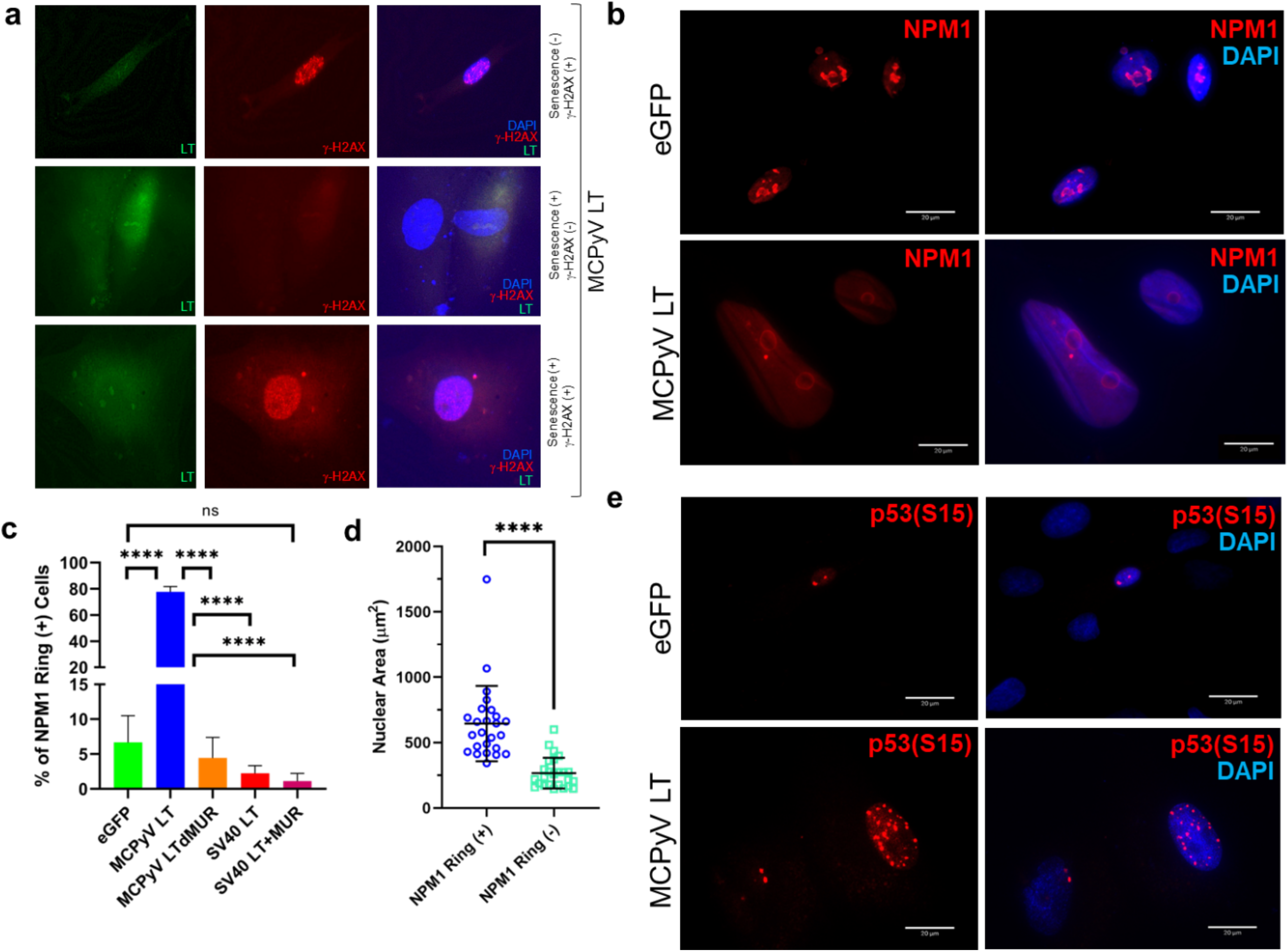
MCPyV LT induces a nucleolar stress response (NSR). (**a**) MCPyV LT-induced senescence is DDR-independent. Immunofluorescent staining for γ-H2AX, a DNA damage marker, in senescent and nonsenescent MCPyV LT positive cells. (**b**) MCPyV LT reorganizes the nucleolus. MCPyV LT induced a ring-like perinucleolar distribution of NPM1 within the nucleolus, a characteristic of canonical nucleolar stress. Immunofluorescent analysis of NPM1 is shown (red). Scale bars = 20 µm. (**c**) MCPyV LT uniquely alters NPM1 structure. The percentage of NPM1 nucleolar ring positive cells for the empty vector and each LT construct was recorded. Statistical significance was determined using the one-way ANOVA test (**** p < 0.0001, ns = not significant). Standard error bars represent mean value with standard error, n = 30 cells with 3 replicates. (**d**) Cells with NPM1 reorganization induced by MCPyV LT display senescent phenotypes. Nuclear size (µm^2^) of MCPyV LT-expressing cells with or without NPM1 nucleolar rings. Unpaired student t test (**** p < 0.0001, ns = not significant). Standard error bars represent mean value with standard deviation, n = 25 cells per condition for 3 replicates. (**e**) MCPyV LT-induced NSR activates p53. Phospho-p53 (S15) detected in empty vector or MCPyV LT-expressing cells.

The nucleolar stress response is another mechanism that can induce senescence through p53 activation^30^. This response involves translocation and reorganization of nucleolar proteins, such as nucleophosmin 1 (NPM1)^31,32^, to the periphery of the nucleolus or into the nucleoplasm, subsequently activating the p53/p21 pathway, and inhibiting ribosomal RNA (rRNA) synthesis^31^. Senescent MCPyV LT-expressing cells displayed a distinct ring-like nucleolar NPM1 distribution that can be triggered in response to nucleolar stress (Figure 2b)^32–34^. In contrast, empty vector, SV40 LTs, MCPyV LTdMUR-expressing cells exhibited a normal speckled NPM1 staining pattern typically found in unstressed cells (Figure S3a). Approximately 80% of MCPyV LT positive cells formed NPM1 nucleolar rings (Figure 2c) and had an enlarged nuclear size (>500 μm^2^) (Figure 2d) comparable to the nuclear size of senescent cells seen in Figure 1c. These results show that MCPyV LT induces a nucleolar stress response.

Phosphorylation of p53 at serine 15 (S15), a prime event prior to p21 activation, can be induced by the NSR. S15 phosphorylation of p53 was noted in MCPyV LT-positive cells marked by the staining of numerous foci in the nucleus (Figure 2e), but it was substantially reduced in MCPyV LTdMUR (Figure S3b). SV40 LT constructs exemplified phospho-p53 staining (Figure S3b) but did not induce p21 expression as shown in Figure 1f and Figure S1a as previously reported in U2OS cells^13^. Taken together, these results indicate that MCPyV LT-mediated senescence is dependent upon the NSR, leading to activation of the p53/p21 pathway.

### MCPyV LT-induced senescence is p21-dependent

Because p21 has a key role in cellular senescence, we hypothesized that the loss of p21 expression would block MCPyV LT-induced senescence. To determine whether MCPyV LT-mediated modulation of p53/p21 is required for the onset of senescence, knockdown of p21 was performed by short hairpin RNA (shRNA) transduction in BJ-hTERT cells expressing MCPyV LT using two shRNA lentiviral constructs, shp21.1 and shp21.2. Transduction with scrambled shRNA (Scr) was used as a control. Knockdown of p21 was confirmed at the mRNA and protein levels in MCPyV LT-expressing cells by RT-qPCR (Figure 3a) and immunoblotting (Figure 3b), respectively. Knockdown using shp21.1 drastically decreased (∼95%) p21 mRNA and protein levels, while knockdown with shp21.2 reduced p21 mRNA expression to a lesser degree (∼40%) and did not decrease p21 protein levels. Dual knockdown with shp21.1 and shp21.2 (shp21.1/2) reduced p21 to similar levels to shp21.1 knockdown alone. Since it was the most potent at knocking down p21, shp21.1 construct (p21KD) was used for the remaining experiments (Figures 4–6). Knockdown of p21 after lentiviral transduction of MCPyV LT reversed cell cycle arrest, as shown by decreased cyclin E and increased pHH3 expression (Figure 3b). Moreover, p21 knockdown cells no longer exhibited an enlarged and flattened morphology compared to the MCPyV LT-expressing cells transduced with scrambled shRNA control (Figure 3c). The nuclear area of MCPyV LT-expressing cells was significantly reduced from ∼580 µm^2^ to ∼190 µm^2^, approximately the nuclear size of eGFP control cells (Figure 3d). SA-β-gal staining confirmed that p21 knockdown largely inhibited MCPyV LT-mediated senescence (Figure 3e, Figure S4), concluding that MCPyV LT-induced senescence is p21 dependent.

**Figure 3:**
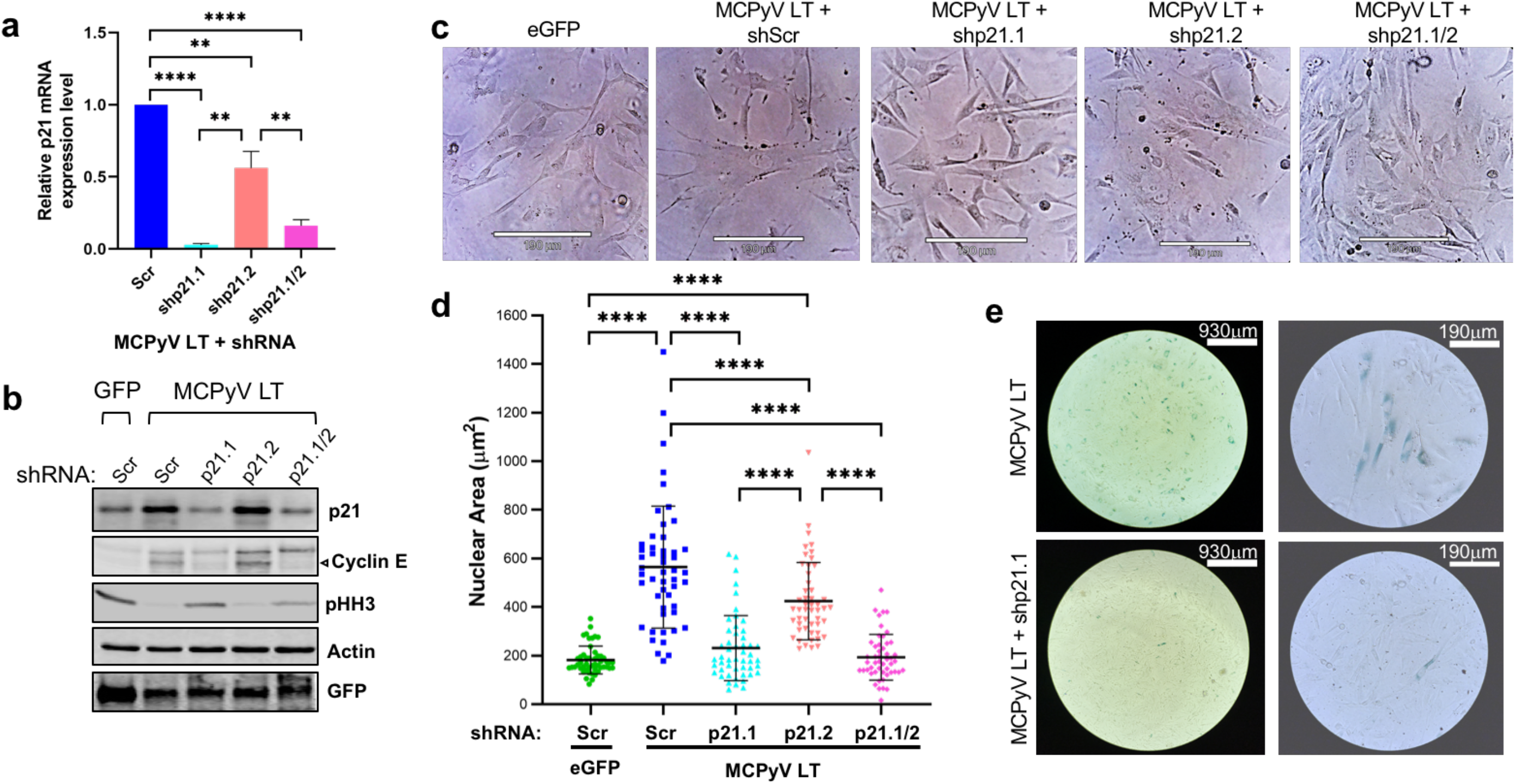
MCPyV LT induces p21-dependent cellular senescence. (**a**) p21 mRNA levels determined after p21 knockdown in MCPyV LT positive cells by RT-qPCR. BJ-hTERT cells were transduced with either empty vector or MCPyV LT and selected with puromycin. At 14 days post transduction, MCPyV LT-expressing cells were transduced with either scrambled (Scr) shRNA or shp21.1, shp21.2, or both (shp21.1/2) for p21 knockdown and then analyzed seven days later (Day 21). Statistical significance was determined using the one-way ANOVA test (** p < 0.01, **** p < 0.0001). Standard error bars represent mean value with standard error, n = 3. (**b**) Analysis of p21 knockdown efficiency by immunoblotting. Both p21 and cyclin E protein levels activated in MCPyV LT-expressing cells were reduced by shRNA transduction. The arrow indicates the major ∼50-kDa band of cyclin E. Histone H3 phosphorylation at Ser10 (pHH3), a mitotic marker, is downregulated in MCPyV LT-positive cells with p21 activation, signifying G2 cell cycle arrest. Actin was used as a loading control. (**c**) Senescent morphological changes in MCPyV LT-expressing cells is reversed after p21 knockdown. Brightfield images of empty vector or MCPyV LT-expressing cells are shown after knockdown. Scale Bar = 190 μm (**d**) Abnormal nuclear size change in MCPyV LT-expressing cells is reversed after p21 knockdown. Statistical significance was determined using the one-way ANOVA test. Standard error bars represent mean value with standard deviation, n = 50 cells. (**e**) p21 knockdown reverses senescence. SA-β-gal senescence assay in MCPyV LT-expressing cells was performed after p21 knockdown.

**Figure 4:**
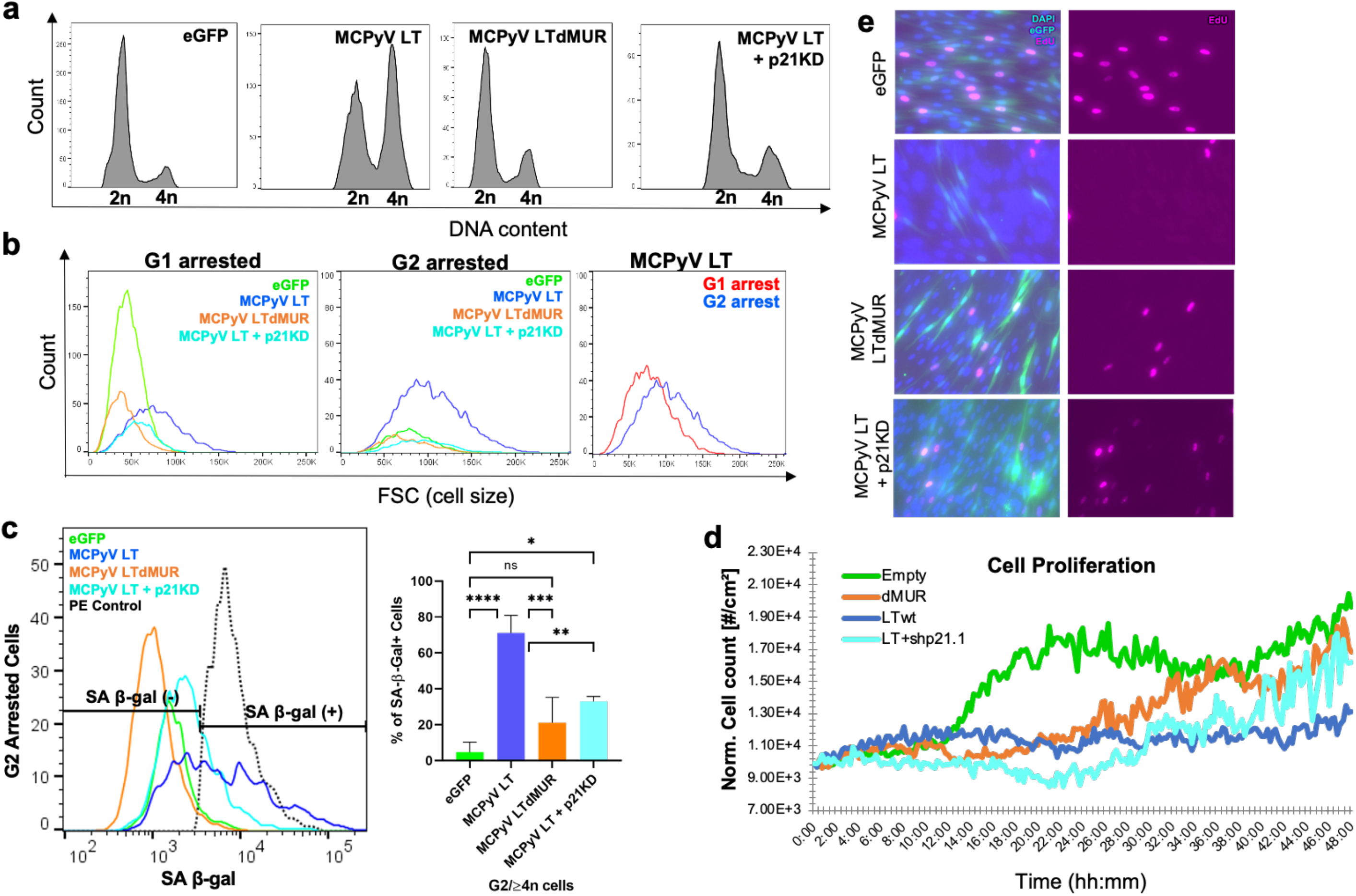
MCPyV LT leads to prolonged G2 cell cycle arrest. (**a**) MCPyV LT-expressing cells accumulate mainly in G2 phase. Cell cycle analysis of eGFP empty vector, MCPyV LT, MCPyV LTdMUR, and MCPyV LT + p21KD (shp21.1) using flow cytometry. Hoechst 33342 was used to stain DNA. 10,000 events were analyzed for each sample, n = 3. (**b**) Histogram of forward scatter (FSC) comparisons of G1- and G2-arrested cells. G1 and G2 cells in MCPyV LT-expressing (GFP^+^) cells were gated and analyzed for cell size using forward scatter (FSC) parameter. Cellular size of G2-arrested cells in MCPyV LT-expressing cells is larger than G1-arrested cells. (**c**) MCPyV LT-induced senescent cells are arrested mainly in G2. Histogram of SA-β-gal staining (PE) on G2 arrested cells (left). Quantification of G2-arrested/SA-β-Gal+ cells by flow cytometry (right). Black dashed line represents PE positive control. Statistical significance was determined using the one-way ANOVA test (* p < 0.05, ** p < 0.01, *** p < 0.001, **** p < 0.0001, ns = not significant). Standard error bars represent mean value with standard error. 10,000 events were analyzed for each sample, n = 3. (**d**) MCPyV LT restricts cell proliferation by inducing senescence. Noninvasive live cell imaging of transduced BJ-hTERT cells was conducted using a phase holographic imaging system (Holomonitor M4) to measure cell proliferation rates. (**e**) MCPyV LT inhibits cell proliferation. Cell proliferation assay. EdU incorporation (pink) was measured by fluorescence microscopy.

### Senescent MCPyV LT-expressing cells are arrested in G2 phase

p21-induced senescence commonly leads to G1 cell cycle arrest ^1^. To verify the cell cycle status and further characterize the growth arrest modulated by MCPyV LT, cell cycle and cell proliferation analyses were performed. Flow cytometric analysis showed that MCPyV LT induced G2 cell cycle arrest, as seen by enrichment of cells in G2 phase (Figure 4a). In contrast, all other LT constructs displayed minimal accumulation of cells in G2 phase (Figure S5a, S5b). Further analysis of MCPyV LT G1- and G2-arrested cells revealed that these cells were larger than the control and MCPyV LTdMUR-expressing cells, suggesting that these growth arrested cells were likely senescent (Figure 4b). G2-arrested cells were larger than their G1-arrested counterparts, implying that G2-arrested cells were senescent. This possibility was confirmed by the observation that approximately 70% of MCPyV LT G2-arrested cells were positive for SA-β-gal. In contrast, only ∼20% of MCPyV LTdMUR G2 cells were positive for SA-β-gal (Figure 4c). As expected, knockdown of p21 in MCPyV LT-expressing cells induced cell cycle reentry (Figure 4a), reduced cell size to eGFP control levels (Figure 4b), and decreased SA-β-gal expression to near MCPyV LTdMUR levels (Figure 4c).

MCPyV LT expression has been shown to reduce cell proliferation through either the C-terminus or the MUR domain^13–15^. To identify if cellular senescence was a potential cause of growth inhibition though the MUR domain, non-invasive live imaging and cell counting was conducted using a HoloMonitor (PHI Lab)^35^. MCPyV LT-expressing cells proliferated slower than their MCPyV LTdMUR counterparts (Figure 4d). Additionally, incorporation of 5-ethynyl-2’-deoxyuridine (EdU) by MCPyV LT-expressing cells was markedly reduced compared to the MCPyV LTdMUR mutant (Figure 4e, Figure S6). Knockdown of p21 in MCPyV LT-expressing cells reversed growth inhibition induced by MCPyV LT expression (Figure 4d, 4e). Although p21-dependent senescence often leads to G1 cell cycle arrest^1^, our results indicate that MCPyV LT-mediated p21 activation mainly facilitated G2-arrested cellular senescence, negatively regulating cell growth.

### MCPyV LT-induced senescence alters host gene expression profile

Senescent cells are characterized by the enhanced secretion of cytokines known as SASPs^1^. To further characterize MCPyV LT-induced senescence, RNA sequencing was performed (Figure 5). Multidimensional analysis indicated that replicates were transcriptionally similar and clustered together (Figure S7a). Of note, cells expressing eGFP or dMUR had similar transcriptomes, further demonstrating the importance of the MUR domain on LT function (Figure S7a, S7b). Additionally, p21 knockdown (p21KD) drastically altered gene expression. Volcano plots illustrated many differentially expressed genes between MCPyV LT- and LTdMUR-expressing cells and in MCPyV LT-expressing cells with p21KD (Figure S7c). A heatmap containing the top 100 differentially expressed genes (DEG) was generated comparing eGFP, MCPyV LT, MCPyV LTdMUR, and MCPyV LT+p21KD (Figure 5a). Numerous SASPs including CXCL1, CXCL2, CXCL5, IL1a, IL1b, IL6, IL8, SAA1, SAA2 were identified on this heatmap and MCPyV LT led to a modest upregulation of some of these SASPs compared to the MCPyV LTdMUR and eGFP control. Unexpectedly, p21KD led to a dramatic increase in expression of inflammatory cytokines, potentially due to cell death induced by p21KD after lentiviral transduction.

**Figure 5:**
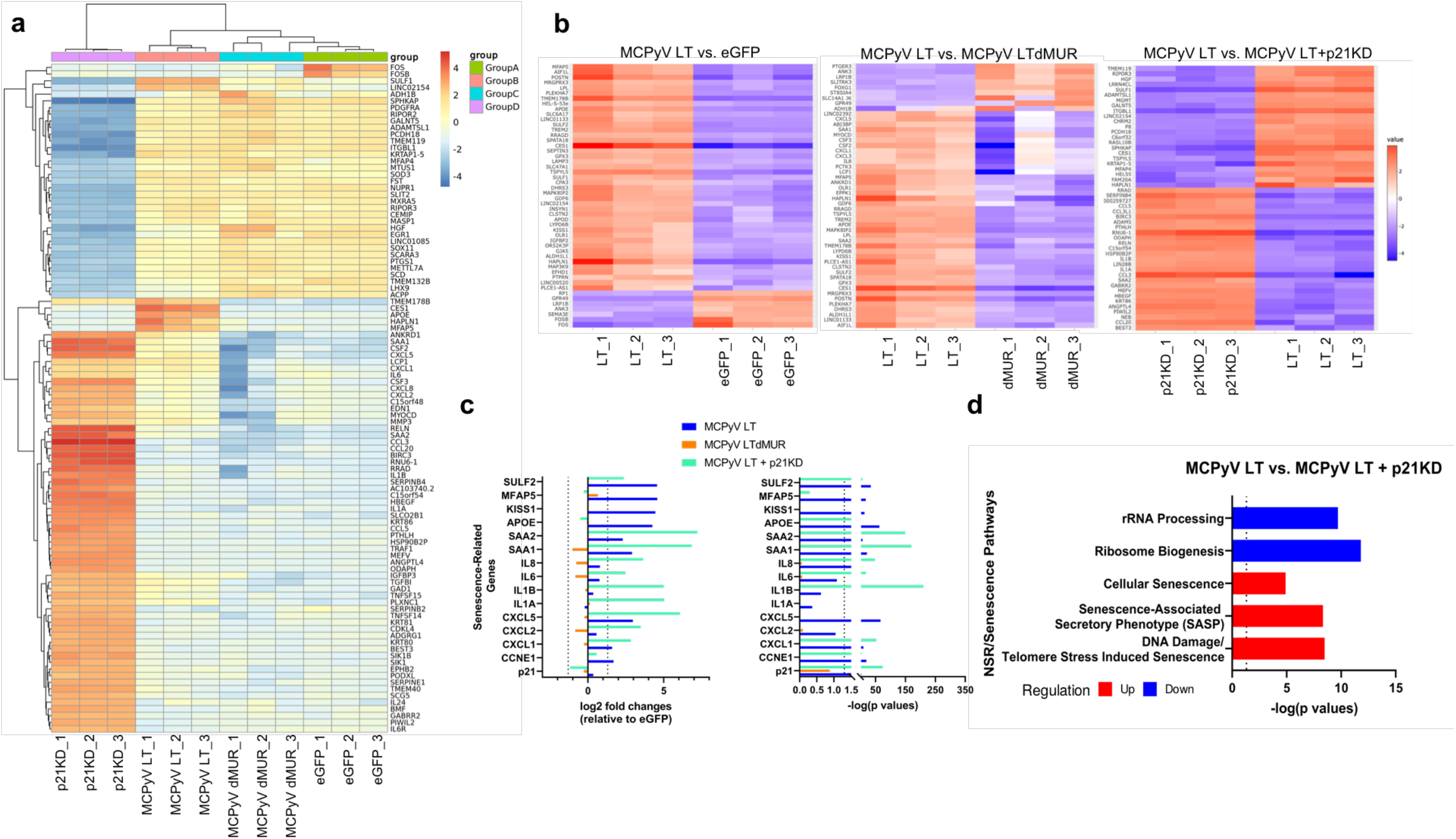
MCPyV LT-expressing cells are enriched in senescence-related genes. (**a**) Heatmap comparing top 100 differentially expressed genes comparing eGFP, MCPyV LT, MCPyV LTdMUR, MCPyV LT+p21KD. (**b**) Heatmaps comparing top 50 differentially expressed genes between MCPyV LT:eGFP (left), MCPyV LT:MCPyV LTdMUR (middle), MCPyV LT:MCPyV LT+p21KD (right). Heatmaps were generated using iDEP. (**c**) SASPs expression. Expression of senescence-related genes compared to eGFP empty vector cells. Log_2_ fold changes (left), positive values represent upregulated genes compared to eGFP cells, negative values represent downregulated genes compared to eGFP cells. Vertical dashed lines represent +/- 1.5 log_2_ fold change with a false discovery rate (FDR) of 0.1. −log(p values) of senescence genes (right). Dotted line represents p value of 0.05, −log(0.05) = 1.3. Abbreviations – SULF2: Sulfatase 2, MFAP5: Microfibril associated protein 5, KISS1: Kisspeptin 1, APOE: Apolipoprotein E, SAA: Serum Amyloid A1 and A2, IL: Interleukin, CCNE1: Cyclin E1. (**d**) Senescence pathways enriched in MCPyV LT compared to MCPyV LT+p21KD. Dotted lines represent p value of 0.05, −log(0.05) = 1.3 and p values to the right of the dotted line < 0.05 (significant).

To further investigate the changes in gene expression due to MCPyV LT, the MUR domain, and p21-dependent senescence, we generated heatmaps of the top 50 differentially expressed genes (DEG) comparing MCPyV LT to eGFP, MCPyV LTdMUR, and p21KD, respectively (Figure 5b). SASPs and senescence genes commonly found as a top DEG were examined. Log_2_ fold change of genes and their corresponding p values compared to eGFP were plotted (Figure 5c). Cyclin E transcripts were significantly upregulated compared to empty vector and the dMUR mutant (p value<0.05). Significant SASPs that were upregulated in MCPyV LT-expressing cells compared to eGFP and dMUR mutant cells included SAA1, SAA2, CXCL1, and CXCL5. Pathway analysis indicated that MCPyV LT-expressing cells were enriched in senescence-related pathways and downregulated in rRNA and ribosomal metabolic pathways compared to p21KD cells (Figure 5d), reinforcing the conclusion that MCPyV LT promotes NSR to activate p21-mediated senescence^36^. Genes found in the enriched senescent pathways included numerous histone variants (Supplementary dataset), which have been previously associated with senescence^37^. Some of the expressed aging/senescence-related genes found highly upregulated in MCPyV LT^+^ cells (SULF2^38^, MFAP5^39^, KISS1^40^, and APOE^41,42^) were inversely regulated by p21KD (Figure 5c, 5d).

### MCPyV LT-mediated senescence maintains MCPyV genome

As senescent cells are characterized by a stable cell cycle arrest and are resistant to apoptosis ^4^, we asked if MCPyV could exploit cellular senescence to establish persistent viral infection. To test the hypothesis that senescence is required for longevity of the MCPyV infection and do so in the context of an actual MCPyV molecular clone, we transfected BJ-hTERT cells with a full length MCPyV genome (pMCPyV-MC) where the early genes were tagged with eGFP (Figure 6a). Senescent MCPyV cells were present at 21 days post transfection, whereas knockdown with p21 greatly abolished SA-β-gal staining (GFP^+^/SA-β-Gal^+^) as determined by both flow cytometry and microscopic analysis (Figure 6a, 6b and Figure S8a).

**Figure 6:**
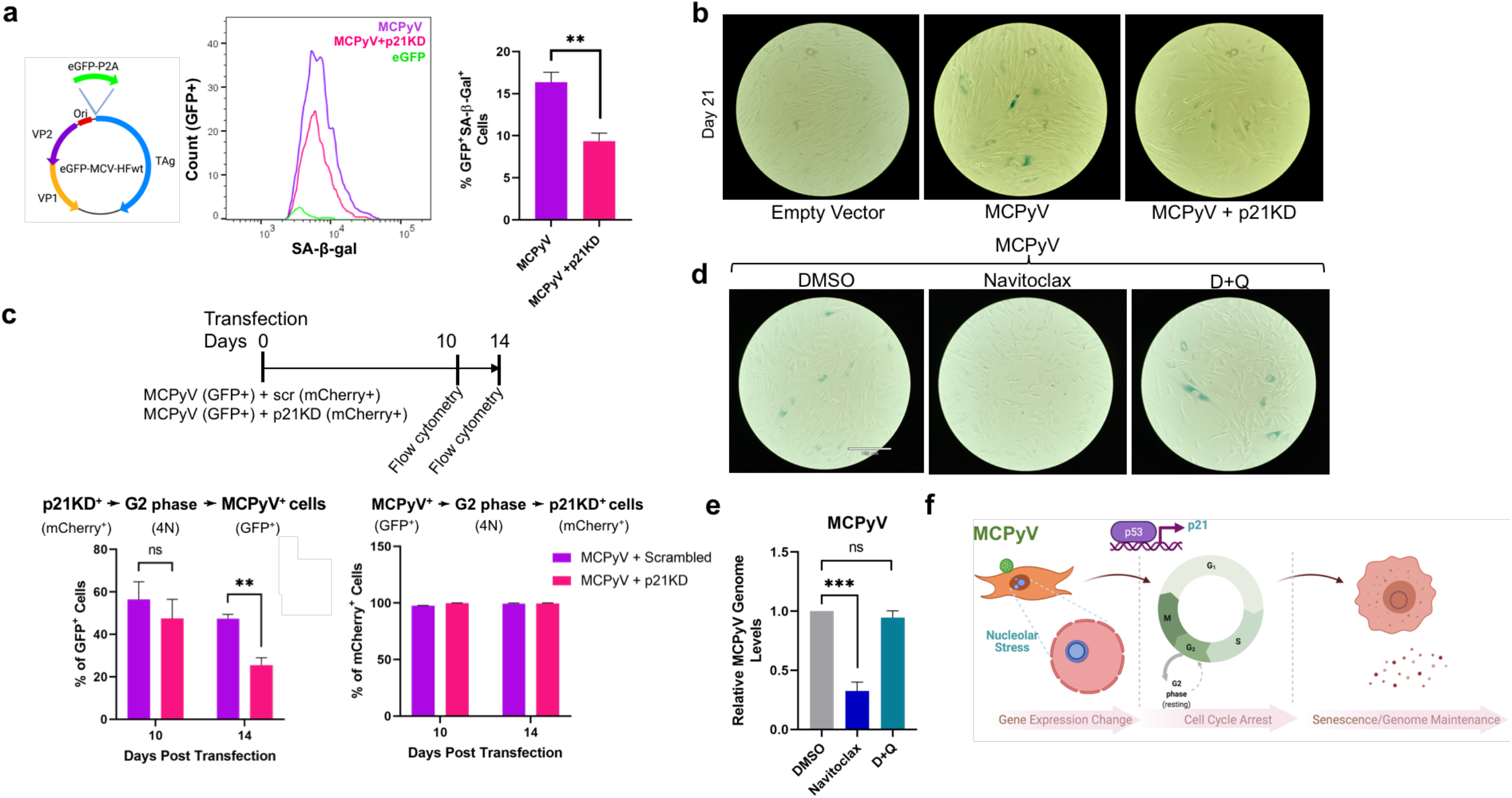
MCPyV LT-induced senescence is required for viral genome maintenance. (**a**) p21 knockdown decreases the percentage of senescent (SA-β-gal^+^) cells. Schematic representing whole MCPyV genome construct (pMCPyV-MC) encoding origin (Ori), VP1, VP2, T antigens (TAg), and eGFP-P2A (left). BJ-hTERT cells were reversely transfected with the MCPyV genome and then transduced with shp21.1 for p21 knockdown 14 days post transfection. At 21 days post transfection, MCPyV transfected cells were subjected to a senescence assay. Histogram representing GFP^+^/SA-β-gal^+^ (MCPyV^+^, senescent cells) for transfected MCPyV or MCPyV+p21KD cells at 7-days post p21 knockdown (middle). Percentage of GFP^+^/SA-β-gal^+^ in population (right). Statistical significance was determined using an unpaired student t test (** p < 0.01). Standard error bars represent mean value with standard error, n = 3, 10,000 cells recorded per replicate. (**b**) MCPyV-induced senescence is p21 dependent. Colorimetric SA-β-gal staining of MCPyV transfected cells at 21 days post transfection. (**c**), LT-induced cellular senescence maintains MCPyV genome. BJ-hTERT cells were cotransfected with MCPyV (GFP+) and mCherry-shScr or mCherry-shp21.1 (p21KD, mCherry+). Cells were harvested at stated days post transfection and were analyzed by flow cytometry. Transfected cells were gated for mCherry^+^ (p21KD) cells and then examined for G2-arrested GFP^+^ cells (MCPyV^+^) (left). Cells were first gated for GFP^+^ cells and then G2 arrested cells that were also mCherry^+^ to ensure cotransfection of both MCPyV and shRNA constructs (right). Unpaired student t test (** p < 0.01, ns = not significant). Standard error bars represent mean value with standard error, n = 3. (**d-e**) Senolytic treatment selectively clears LT-induced senescent cells and reduces MCPyV genome levels. At 14 days post transfection, empty vector- or MCPyV-transfected cells were treated with navitoclax (250 nM), dasatinib+quercetin (D+Q, 250 nM, 25 nM dasatinib + 250nM quercetin), or DMSO for two days and then subjected to SA-β-gal assay (**d**) or qPCR (**e**). Scale bar = 180 μm. One way ANOVA (*** p < 0.001). Standard error bars represent mean value with standard error, n = 3. (**f**) MCPyV induces senescence to sustain viral genome maintenance. MCPyV LT activates p53 and subsequently p21 through nucleolar stress responses. Activation of p21 induces G2 cell cycle arrest, leading to changes in growth arrest and senescent gene expression and promotion of cellular senescence. Cellular senescence establishes MCPyV infection in fibroblast cells, enabling it to extend its lifecycle within the host. Figure created with BioRender.com.

To determine if senescence is required for MCPyV genome maintenance, we reversely cotransfected BJ-hTERT cells with the MCPyV genome and mCherry-Puro shScr or mCherry-Puro shp21.1. Transfected cells were harvested at 2-, 4-, and 7-days post transfection and then subjected to RT-qPCR and qPCR to confirm knockdown of p21 transcripts and observe viral genome levels over time respectively (Figure S8b). Cells were initially transfected with shp21.1 to inhibit senescence, as we sought to determine how loss of senescence impacts MCPyV long-term persistence. Though not significant, at days 4 and 7 post transfection, p21 knockdown reduced MCPyV genome levels by 50% compared to scrambled-transfected cells. We also observed MCPyV genome maintenance using flow cytometry by detecting GFP levels over time (Figure 6c). Since MCPyV LT induced G2 cell cycle arrest (Figure 4a), we specifically examined the G2 population. Cells were gated on GFP and mCherry (GFP^+^mCherry^+^) to ensure only cells that had taken up both the MCPyV genome and shRNA plasmid were examined (Figure 6c). At day 14 post transfection, p21 knockdown reduced the percentage of GFP+ cells by ∼25% compared to scrambled control cells.

Previous studies have identified senolytics as drugs that can selectively clear senescent cells to reduce disease morbidity ^1,10,43^. Therefore, we investigated whether or not senolytic treatment could potentially act as a therapeutic agent to prevent MCC. Fourteen days post-transfection, cells were treated with navitoclax, dasatinib+quercetin (D+Q), or DMSO for two days. To determine if these senolytics could reduce the senescent cell burden, a SA-β-Gal assay was conducted. Navitoclax treatment ^44^ could markedly reduce SA-β-Gal staining, whereas D+Q treatment did not show a significant change in SA-β-Gal staining (Figure S9, Figure 6d). Moreover, only navitoclax could reduce viral genomes levels, decreasing MCPyV DNA levels by ∼70% (Figure 6e).

## Discussion

Taken together, our results showed that MCPyV infection, mainly MCPyV LT expression, can activate a nucleolar stress response in human fibroblast cells. This in turn activated p53 transcriptional activity, inducing p21 expression. p21 activation subsequently led to G2 cell cycle arrest, promoting cellular senescence. MCPyV was able to exploit these senescent cells by maintaining its viral genome for long-term survival (Figure 6f). MCPyV-induced senescence was recently reported by Siebels et al^45^. Although their MCPyV LT interaction study revealed an interaction with KRAB-associated protein 1 (KAP1), a senescence-related factor^46^ in primary normal human dermal fibroblasts (nHDF), it was not clear if LT expression alone is sufficient to induce host cellular senescence. Of note, Siebels et al. observed neither senescence nor p21 transcriptional activation assessed in primary nHDFs in 2-4 days of LT expression alone. In agreement with that, we did not detect significant morphologic changes by LT transduction in primary nHDF (Figure S11) even 7 days post transduction. As an unexpected high SA-β-gal positive background was observed in early passages of non-immortalized skin fibroblast cells (HFF-1 and BJ), including nHDF (Figure S11), we used BJ-hTERT, a skin fibroblast cell line commonly used to study aging and senescence^47–49^ for all our experiments to determine accurate gene changes in MCPyV LT-induced senescence. Although BJ-hTERT cells express human telomerase reverse transcriptase (hTERT), which is known to inhibit senescence, it has been shown that hTERT has no effect on stress-induced senescence^50^, but only inhibits stress-related apoptosis and necrosis^51^.

In latent viral infection, while the full viral genome is retained in the host cell, its late gene expression is dramatically restricted and only few viral antigens and no viral particles are produced. To qualify as latency, this quiescent state of infection must display persistence and reversibility ^52^. Protein-mediated viral latency is the only known mechanism for human polyomavirus latency ^17^, shown by successful reversibility from a quiescent state to a productive infection that is regulated by MCPyV LT proteostasis^17^. Accompanied by the protein interaction networks^15,17,45,53^, here we show that an additional key nucleolar stress pathway, triggered by MCPyV LT overexpression, plays a pivotal role in persistent viral infection. Although viral replicative machinery has been associated with modulating the nucleolus function, none of the DNA viral oncoproteins have been specifically characterized to trigger changes in nucleolar distribution^36,54^. Canonically, induction of the nucleolar stress response is caused by disruption of ribosomal biogenesis^31^. Though our RNA-Seq data suggests MCPyV LT can downregulate rRNA synthesis, the mechanism of NSR induction by MCPyV LT needs further investigation. Certainly, other stress responses in different cell types such as DDR, shortened telomeres, replicative stresses, and/or apoptosis will combinedly influence the extent of senescence, in early and late stages of the infection process which can be further explored as well. Nonetheless, it is important that both persistency and reversibility are mainly modulated by the MCPyV LT unique domain that is potentially involved in a series of host interactions and regulation of its own proteostasis^15,17,55,56^.

Most of the known human cancer viruses have an ability to antagonize p53 by expressing viral oncoproteins that promote cellular proliferation by abrogating p53-induced cell cycle arrest in response to DNA damage^57^. Human polyomavirus LTs have been known to interact with p53, which immediately blocks DDR and innate immune sensing for host survival in the initial infection process^58^. Unlike other human polyomaviruses, MCPyV LT does not seem to possess an ability to interact and directly inhibit p53^14^, but rather activate p53 phosphorylation and function^13^. This unexpected activation of p21 transcription by p53, uniquely induced by MCPyV LT, may establish a distinct niche of MCPyV as a human cancer-causing virus among human polyomaviruses. Low mutation rates of p53^53,59^ and an intact p53/p21 pathway^60^ in MCCs consistently support our results that the activation of p21-dependent senescence potentially plays a role in MCPyV infection. It should be noted that the LT-induced senescence response in human fibroblasts has been simultaneously evolving over time of passages potentially due to changes in miRNA and gene expression profile^61^, suggesting a variety of gene expression changes can alter and allow host cellular responses to support latent infection periods. Specifically, we observed that late passages of BJ-hTERT cells supported enhanced senescence and reduced cell death induced under LT expression and p21 knockdown conditions compared to early passaged cells. We also noticed a significant level of mitotic cell death as an early event induced by p21 knockdown, although cells recover over time. This p21 knockdown-mediated cell death may be caused by re-entry of senescent cells back into the cell cycle, which enhances activation of DDR as shown by others^62^.

Senescence has been characterized as an irreversible exit of the cell cycle^63^. Given that certain viruses depend on host cell proliferation for viral replication, viral oncogene-mediated cell senescence has been suggested to be an antiviral mechanism for host cells. Our results define a central role of senescence that may contribute to viral oncogenesis that has not been previously described. Despite this cell intrinsic tumor suppression, senescent cells have also been implicated as active contributors to tumorigenesis by extrinsically promoting many hallmarks of cancer, including evasion of the immune system^64^. Through induction of cellular senescence, MCPyV potentially creates a precancerous environment in which it can avoid the immune system that enhances cell survival and viral persistence. The senolytic drug navitoclax is the B-cell lymphoma (Bcl-2) anti-apoptotic family protein inhibitor. Early studies on navitoclax have shown promising results to suppress MCC growth^65,66^, suggesting the Bcl-2 pathway as a potential therapeutic target to prevent MCPyV infection and MCC development. Though whether latent viral infection can ultimately lead to oncogenesis by inducing host senescence and other stress environments would need further investigation, such cellular responses surely contribute to maintaining cellular homeostasis during virus infection. Our results uncover the mechanism of a host stress response regulating human polyomavirus genome maintenance in persistent viral infection, which may lead to the targeted intervention for MCC.

## Methods

### Cell culture and plasmids

BJ-hTERT, BJ (ATCC) and 293FT cells (Invitrogen) were cultured in Dulbecco’s modified Eagle’s medium (DMEM) with 10% premium grade fetal bovine serum (FBS, Seradigm). HFF-1 cells (ATCC) were grown in 15% FBS in DMEM. Primary neonatal nHDF cells (CC-2509, Lonza) were supplemented with FGM-2 growth medium-2 BulletKit™ (Lonza). Plasmids constructs encoding codon-optimized cDNA for MCPyV LT, MCPyV LTdMUR, SV40 LT, and SV40 LT+MUR were cloned into pLVX-GFP-P2A empty vectors as previously described ^15^. Enhanced GFP (eGFP) was cloned into MCPyV genome (pMCPyV-MC) to visualize MCPyV early gene expression (Figure S10).

### Lentiviral transduction and p21 knockdown

For lentivirus production, 293FT cells (Invitrogen) were co-transfected with lentiviral psPAX2 packaging, pMD2.G envelop, and pLVX or pLKO.1 transfer plasmids (Addgene) using jetOPTIMUS according to the manufacturer’s instructions, except the amounts of each plasmid was increased for optimal lentiviral production. BJ-hTERT cells were transduced with lentivirus encoding pLVX-GFP-P2A empty vector, MCPyV LT, MCPyV LTdMUR, SV40 LT, or SV40 LTMUR in the presence of 6 µg/mL polybrene (Sigma-Aldrich). For stably transduced BJ-hTERT cells, the transfectants were selected and maintained with 3 µg/ml puromycin (Sigma-Aldrich). MCPyV LT-expressing BJ-hTERT cells were grown for 14 days and then transduced with scrambled (Scr) short hairpin RNA (shRNA), shp21.1, shp21.2, or both shp21.1/2 for p21 knockdown analysis. shRNA sequence of p21 (shp21.1:TRCN0000287021, shp21.2:TRCN0000040126) were cloned into lentiviral vector pLKO.1 neo (Addgene#13425) or pLKO.1 mCherry-Puro generated by overlapping PCR using primer pairs (Table S1) using AgeI and EcoRI. After transduction, cells were selected with 500 µg/mL geneticin (G418, Gibco) for two days. Knockdown efficiency was confirmed by reverse transcription-quantitative PCR (RT-qPCR) using p21 primer pairs listed in Table S1 and immunoblots seven days after p21 knockdown (21 days post-transduction). A senescence assay and immunoblot analysis was conducted after at 14 days (without p21 knockdown) or 21 days post transduction (with p21 knockdown).

### Quantitative immunoblotting and antibodies

Stably transduced BJ-hTERT cells (selected with puromycin for 14 days) were lysed with IP buffer (50 mM Tris-HCl (pH 8.0), 150 mM NaCl, 1% TritonX-100, 1 mM PMSF, 1 mM benzamidine) and whole cell lysates were used for immunoblot analysis. Membranes were incubated with primary antibody overnight at 4°C, and then incubated with secondary antibody for 2 hours (h) at room temperature (RT). A quantitative Infrared (IR) imaging system, Odyssey CLX (LI-COR) was used to determine protein expression. Primary antibodies used in this study included phospho-p53 (Ser15) (Cell Signaling, 9284), p21 Waf1/Cip1 (Cell Signaling, 2947), Cyclin E (Santa Cruz Biotechnology, sc-247), Phospho-Histone H3 (Ser10) (Cell Signaling, 53348), GFP (Santa Cruz Biotechnology, sc-9996), and β-Actin (Cell Signaling, 4970). All signals were detected using quantitative Infrared (IR) secondary antibodies (IRDye 800CW goat anti-mouse, 800CW goat anti-rabbit, 680LT goat anti-rabbit IgG, 680LT goat anti-mouse IgG) (LI-COR).

### Immunofluorescence assay

Transduced BJ-hTERT cells were seeded in 6-well plates or 8-well chamber slides (Nunc™ Lab-Tek™). Cells were fixed in 4% paraformaldehyde in PBS (Alfa Aesar) for 10 minutes and then permeabilized in 1% Triton X-100 (Sigma-Aldrich) in PBS for 20 minutes. 3% BSA/0.1 M glycine in PBS was used to block for 1 h. Cells were incubated overnight at 4°C with primary antibodies: γ-H2AX(Ser139) (Cell Signaling, 9718), p21 Waf1/Cip1 (Cell Signaling, 2947), pHH3 (Cell Signaling, 53348), p-p53(S15) (Cell Signaling, 9284), and NPM1 (FC-61991, Life Technologies). Secondary antibody (A11029 or A11036, Alexa Fluor 568-conjugated goat anti-mouse or anti-rabbit IgG (H+L) Highly Cross-Adsorbed, Invitrogen) was used to stain cells for 1 h. Cells were incubated with DAPI (0.1 µg/ml, Thermo Scientific) for 5 minutes and then analyzed by fluorescence imaging using a REVOLVE4 microscope (Echo Laboratories).

### Senescence assay

Detection of SA-β-Gal activity was performed using a colorimetric senescence β-Galactosidase histochemical staining kit (G-Biosciences) or a fluorescent Cell Meter™ Cellular Senescence Activity Assay Kit with Xite™ Red beta-D-galactopyranoside (AAT Bioquest) according to the manufacturer’s instruction. Briefly, the cells were washed with PBS and then fixed with 4% paraformaldehyde in PBS for 15 min. Cells were then permeabilized for 15 minutes with 0.1% Triton in PBS or permeabilization buffer (Invitrogen). After incubation with SA-β-Gal detection solution for 12 h (G-Biosciences) or 45 minutes (AAT Bioquest) at 37ºC, the cells were stained with DAPI (Thermo Scientific) for microscopic or Hoechst 33342 (Invitrogen) for flow cytometric analysis. For LT-transduced cells, senescence assays were performed at 14 days or 21 days post-transduction for samples with or without p21 knockdown respectively. Normal fibroblast cell lines were stained for SA-β-Gal 7 days post-transduction. MCPyV genome-transfected cells were transduced with shp21.1 at 14 days post transfection. At 21 days post transfection, a senescence assay was conducted. Images were captured using a REVOLVE4 fluorescent microscope (Echo Laboratories). All the samples were stained at least in triplicate. 50 cells were counted for each LT construct. Nuclear area was calculated using Echo Pro software (Echo Laboratories). Positive PE signal was detected using the PE-Annexin V (BD Biosciences) or CS&T Research Beads (BD Biosciences, 655050). For PE positive control samples (Annexin-V), BJ-hTERT cells were washed twice with PBS and then heat-killed by incubation at 95ºC for 3 minutes. A 1:1 cell suspension of heat-killed and non-heat-killed cells were stained according to the manufacturer’s instructions. 10,000 events were recorded in triplicate for each sample.

### SASP analysis by RT-qPCR

For RT-qPCR, total RNA was isolated using the Monarch Total RNA Miniprep Kit (New England Biolabs) according to the manufacturer’s instruction. mRNA levels of SASPs (Figure 1e) were measured by RT-qPCR using iTaq Universal One-Step RT-qPCR Kit (Bio-Rad) with primer pairs previously described ^67^. Quantitative analysis was performed using the comparative ΔΔCt method by detecting ribonuclease P (RNase P) and glyceraldehyde 3-phosphate dehydrogenase (GAPDH) as reference genes using primer pairs listed in Table S1. All RT-qPCR experiments included a melting curve analysis to confirm the specificity of the amplicons (95°C for 15 sec, 60°C for 20 sec, and 95°C for 15 sec).

### Cell cycle analysis

BJ-hTERT cells were fixed, permeabilized, and then stained with Hoechst 33342 (Invitrogen) to determine cell cycle stages. Cells were analyzed using an LSR Fortessa flow cytometer (BD Biosciences) and 10,000 events were recorded per replicate. FlowJo (TreeStar, Ashland, OR, USA) software was utilized for data analysis. To examine viable cells, forward scatter (FSC) area versus side scatter (SSC) area density plots were analyzed. Doublets were removed through FSC area versus FSC height gating. Cell cycle analysis was conducted on GFP positive cells to examine only transduced cells.

### EdU incorporation assay

Cells were pulse labelled with EdU (final concentration of 10 µM, Invitrogen) 24 h before fixation. EdU incorporation was detected using the Click-iT Plus EdU Alexa 647 Fluor Imaging Kit (Invitrogen) according to the manufacturer’s instruction. Cells were then stained with DAPI (Thermo Scientific) and visualized with a fluorescence microscope (REVOLVE4, Echo Laboratories).

### Cell proliferation assay

Cells were plated on 6 well, TC-plates (Fisher Scientific) at a confluency of approximately 30% and were incubated with 10% serum. After 24 h, cells were recorded using a phase holographic imaging system (Holomonitor M4 laser microscope, Phase Holographic Imaging, Lund, Sweden)^35^. Images were recorded every 30 minutes for 48 h. The average cell proliferation was quantified using HStudio M4 software (version 2.7.1).

### RNA Sequencing analysis

RNA sequencing and gene reads were aligned to the hg38 human genome. Raw gene counts were used for the integrated Differential Expression and Pathway (iDEP) analysis and were transformed and normalized using iDEP ^68^. All figures and sequence analyses were generated from iDEP or Zymo Research. Genes below 0.5 counts per million (CPM) were excluded and the false-discovery rate (FDR) for differential gene expression was kept below 0.1 (FDR<0.1) with a minimum fold change of 1.5. RNA sequence data are available at NCBI Gene Expression Omnibus (GEO) under the following accession number GSE189291.

### Quantitative PCR Analysis of MCPyV genome

MCPyV genome^69^ with eGFP (pMCPyV-MC, Figure S10) was inserted into a minicircle (MC) parental backbone plasmid pTubb3-MC (Addgene #87112) using SmaI/BsrGI/BstEI sites and isolated according to the manufacturer’s instructions (System Biosciences). BJ-hTERT cells were reversely co-transfected with pMCPyV-MC and pLKO.1 mCherry-Puro shScr (scrambled) or pLKO.1 mCherry-Puro shp21.1 using jetOPTIMUS (Polyplus) following the manufacturer’s instructions. Total DNA was purified using Quick-DNA kit (Zymo Research) at 2, 4, and 7 days post transfection. DNA was digested with DpnI and 20 ng of digested DNA was subjected to qPCR. qPCR was carried out with PowerUp™ SYBR Green Master Mix (Applied Biosystems) using a StepOnePlus™ system (Applied Biosystems) according to the manufacturer’s protocol. Quantitative analysis was performed using the comparative ΔΔCt method by detecting RNase P as a reference gene and three MCPyV detection primer pairs (Table S1). All qPCR experiments included a melting curve analysis to confirm the specificity of the amplicons (95°C for 15 sec, 60°C for 20 sec, and 95°C for 15 sec).

### Flow cytometric analysis of MCPyV genome

BJ-hTERT cells were reverse transfected with pMCPyV-MC and either mCherry-Puro shScr or mCherry-Puro shp21.1 to inhibit senescence and determine its impact on long-term viral maintenance. At days 10 and 14 post transfection, cells were harvested and subjected to flow cytometry to measure GFP and mCherry levels. Detection of mCherry indicated expression of p21 shRNA constructs and observation of GFP indicated presence of MCPyV genome. Cells were first gated on mCherry then followed by gating GFP to ascertain how p21 knockdown affected MCPyV persistence. Gating of GFP then mCherry was used to ensure MCPyV infected cells had taken up the shRNA constructs.

### Senolytic Treatment

pMCPyV-MC or pTubb3-MC (Addgene #87112) transfected cells were maintained for 14 days to allow for senescent phenotypes to be established. Cells were plated onto 24 well or 12 well plates for a SA-β-Gal assay or qPCR analysis respectively and grown for one day. Navitoclax (250 nM), D+Q (250 nM, 25 nM dasatinib plus 250 nM quercetin) (Selleckchem), or DMSO (Sigma-Aldrich) were added to the cells for two days. Senolytic treated cells were then subjected to a SA-β-Gal assay or qPCR to detect viral genome levels, in which 25 ng of undigested DNA was analyzed.

### Statistical analysis

Figures show average values and error bars show the standard error or standard deviation. The one-way analysis of variance (ANOVA) or unpaired student t test was used to determine statistical significance, with p < 0.05 considered significant using GraphPad Prism software (GraphPad Software, Inc., La Jolla, CA, USA). Unless stated, all experiments were tested at least three times in triplicate.

## Acknowledgements

We thank the Department of Microbiology and Immunology and flow cytometry core at Penn State College of Medicine (RRID:SCR_021134) for their support in this project. We also thank Zachary Flegal for assistance on the holographic imaging data analysis and Nnenna Nwogu for preliminary work. This project is funded, in part, under a grant with the Pennsylvania Department of Health using Tobacco CURE Funds. The Department specifically disclaims responsibility for any analyses, interpretations or conclusions.

## Author Contributions

Conceptualization, A.M.P. and H.J.K.; methodology; A.M.P., L.E.O., A.E.L., and H.J.K.; visualization, A.M.P., L.E.O., and H.J.K.; formal analysis, A.M.P., L.E.O., and H.J.K.; writing – original draft, A.M.P. and H.J.K.; writing-review and editing, A.M.P., L.E.O., and H.J.K.; supervision, H.J.K. All authors have read and agreed to the final version of the manuscript.

## Ethics declarations

The authors declare no competing interests.

## Data Availability

All data generated in this study are presented in the paper, supplementary information, supplementary dataset, and NCBI GEO (accession number GSE189291).

