## Supplementary material for "Cellular senescence preserves viral genome maintenance": Table S1, Figure S1-S11

**This PDF file includes:**

Figures S1 to S11
Tables S1

**Other supplementary materials for this manuscript include the following:**

Supplementary Dataset S1

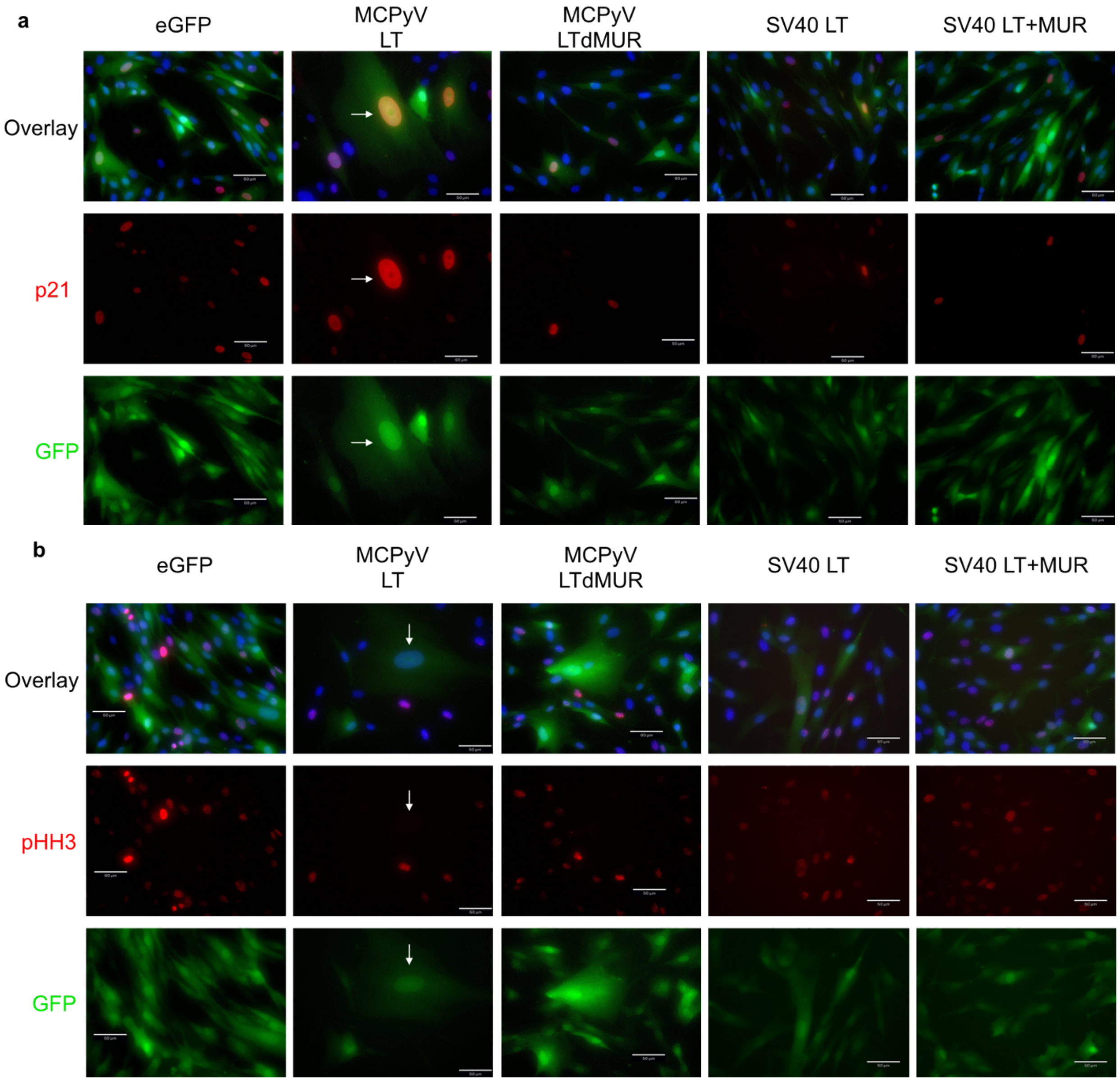


**Fig. S1. MCPyV LT alters expression of growth arrest genes.** Immunofluorescent analysis of p21 (red) **(a)** and pHH3 (phospho-histone H3, red) **(b)** in cells stably expressing eGFP, MCPyV LT, MCPyV LTdMUR, SV40 LT, or SV40 LT+MUR. Enlarged MCPyV LT senescent cells exhibited high p21 expression and low pHH3 levels. SV40 LT constructs did not induce p21 expression. White arrows denote large senescent cells. Scale bar = 60 µm.

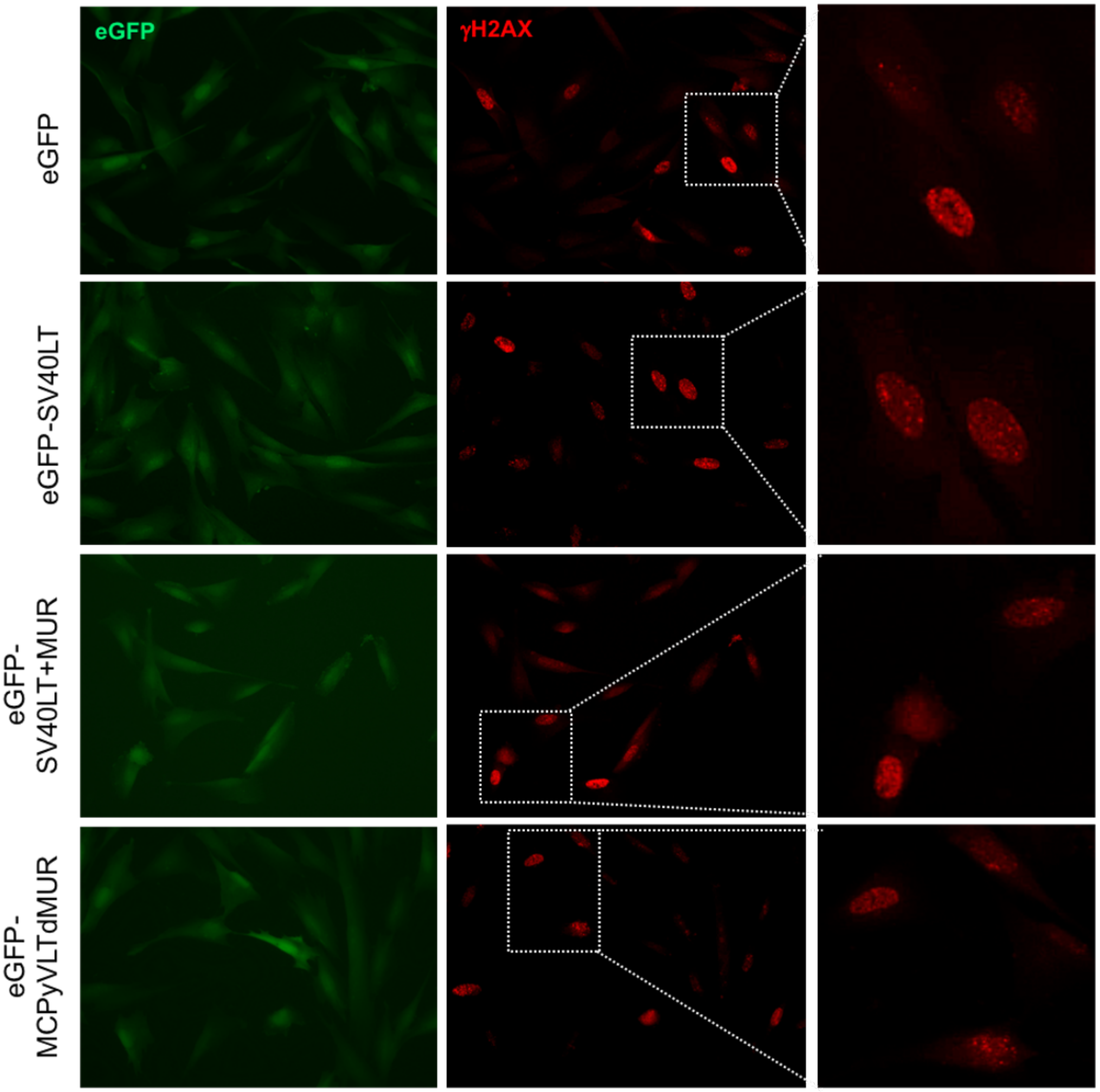


**Fig. S2. Immunofluorescent analysis of γ-H2AX.** Representative immunofluorescence images of nuclear γ-H2AX in eGFP, SV40LT, SV40LT+MUR and MCPyV LTdMUR-expressing cells are shown. Immunofluorescent staining of γ-H2AX (red) shows DNA damage response. Dashed outlines provide higher resolution imaging.

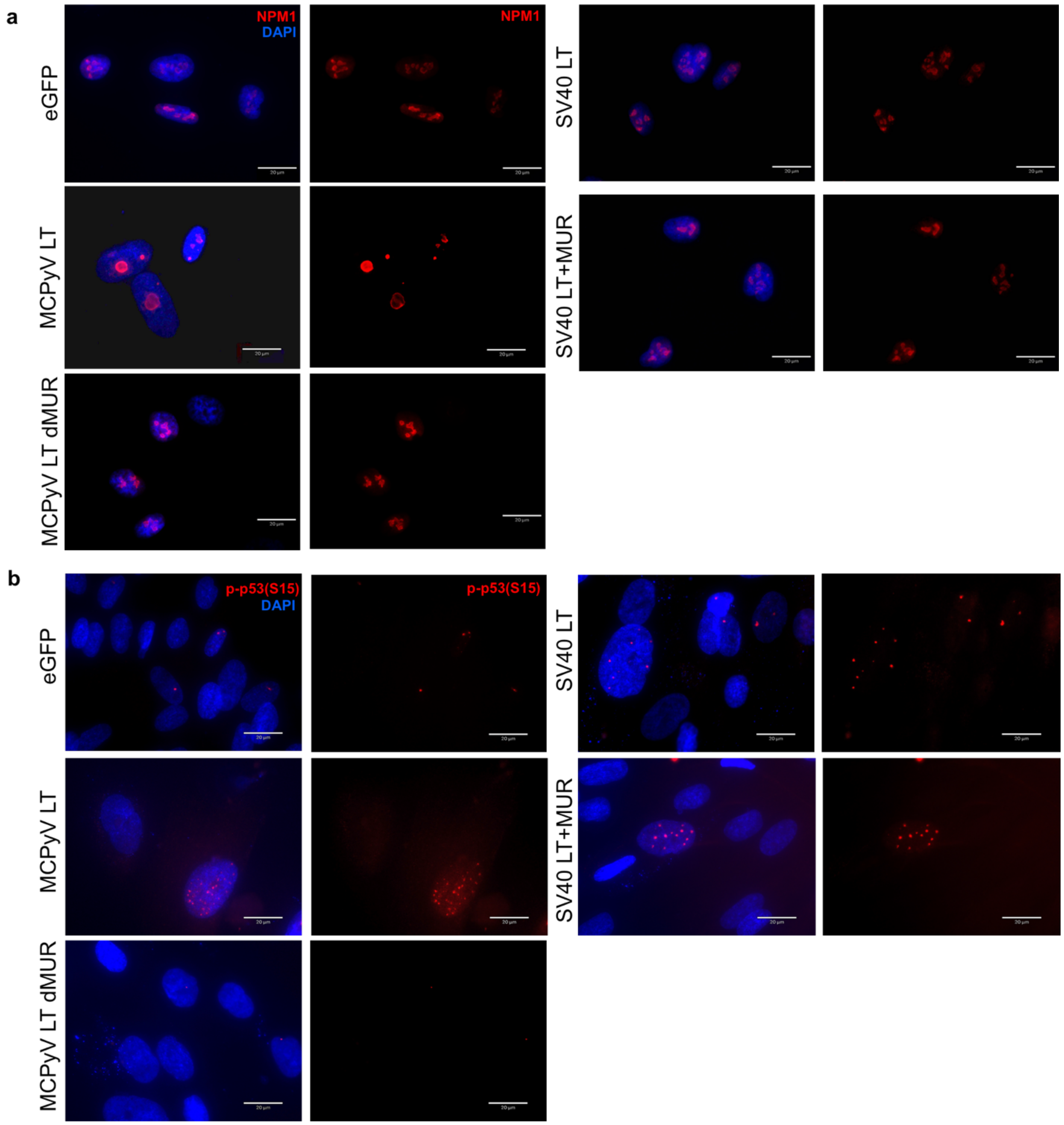


**Fig. S3. MCPyV LT induces a p53-dependent nucleolar stress response. (a)** Senescent MCPyV LT-expressing cells displayed a distinct ring-like nucleolar NPM1 distribution that can be triggered in response to nucleolar stress, while empty vector and MCPyV LTdMUR-expressing cells exhibited a normal speckled NPM1 staining pattern typically found in unstressed cells. **(b)** Phosphorylation of p53 at serine 15 (S15), a prime event prior to p21 activation, can be induced by MCPyV LT expression. S15 phosphorylation of p53 was noted in MCPyV LT-positive cells marked by the staining of numerous foci in the nucleus, but p53 phosphorylation was substantially reduced in cells-expressing MCPyV LTdMUR. SV40 LT constructs exemplified phospho-p53 staining. Scale bar = 20 µm.

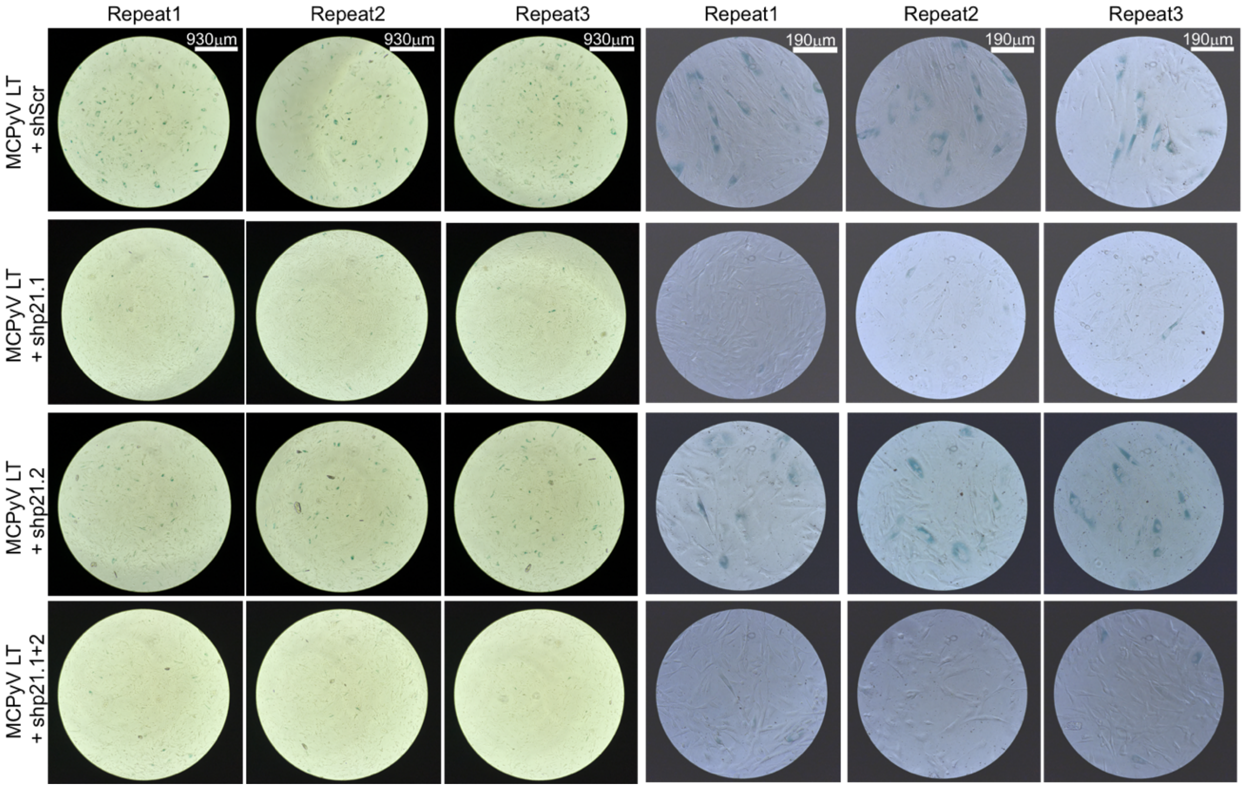


**Fig. S4. MCPyV LT induces p21-dependent senescence.** Knockdown of p21 was performed by short hairpin RNA (shRNA) transduction in BJ-hTERT cells expressing MCPyV LT using two shRNA lentiviral constructs, shp21.1, shp21.2. Transduction with scrambled shRNA (Scr) was used as a control. p21 knockdown largely inhibited MCPyV LT-mediated senescence.

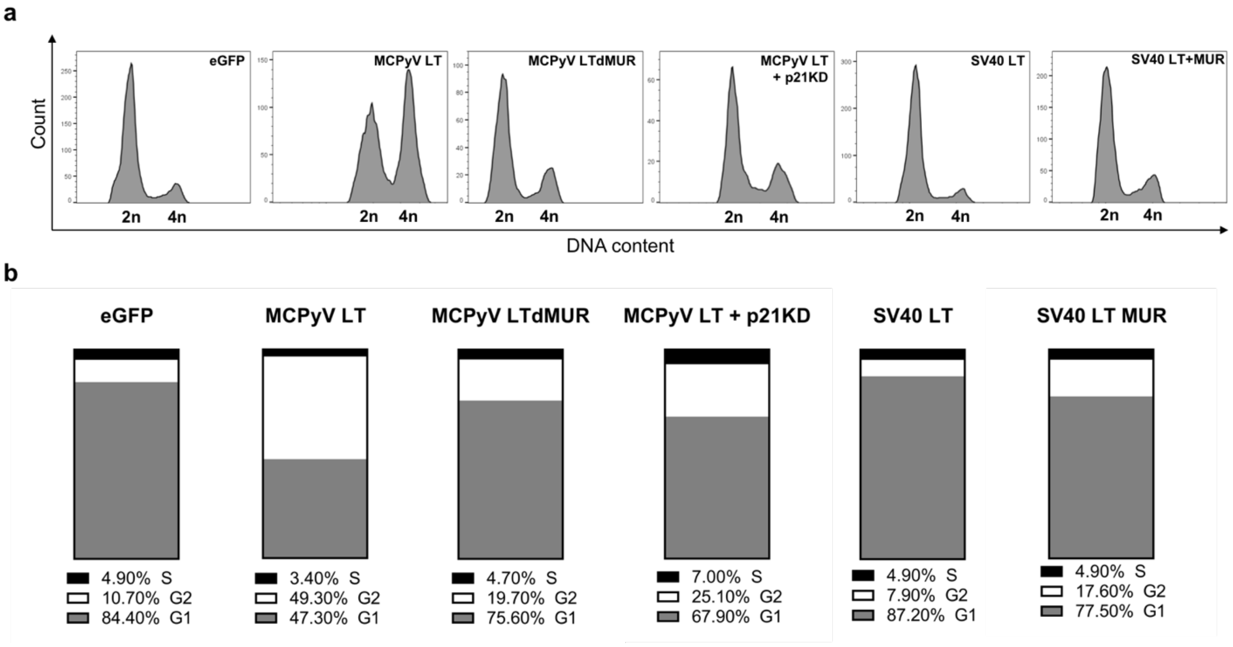


**Fig. S5. MCPyV LT expression induces G2 cell cycle arrest. (a)** Cell cycle analysis using flow cytometry. BJ-hTERT cells-expressing LTs were stained with Hoechst 33342 dye. Cell cycle profiles were denoted as 2n (G1) and 4n (G2). **(b)** Average percentage of cells in G1, S, and G2 phases from three independent experiments is shown. Most of MCPyV LT-expressing cells were in G2 phase. Knockdown of p21 in MCPyV LT-expressing cells allowed for cell cycle reentry.

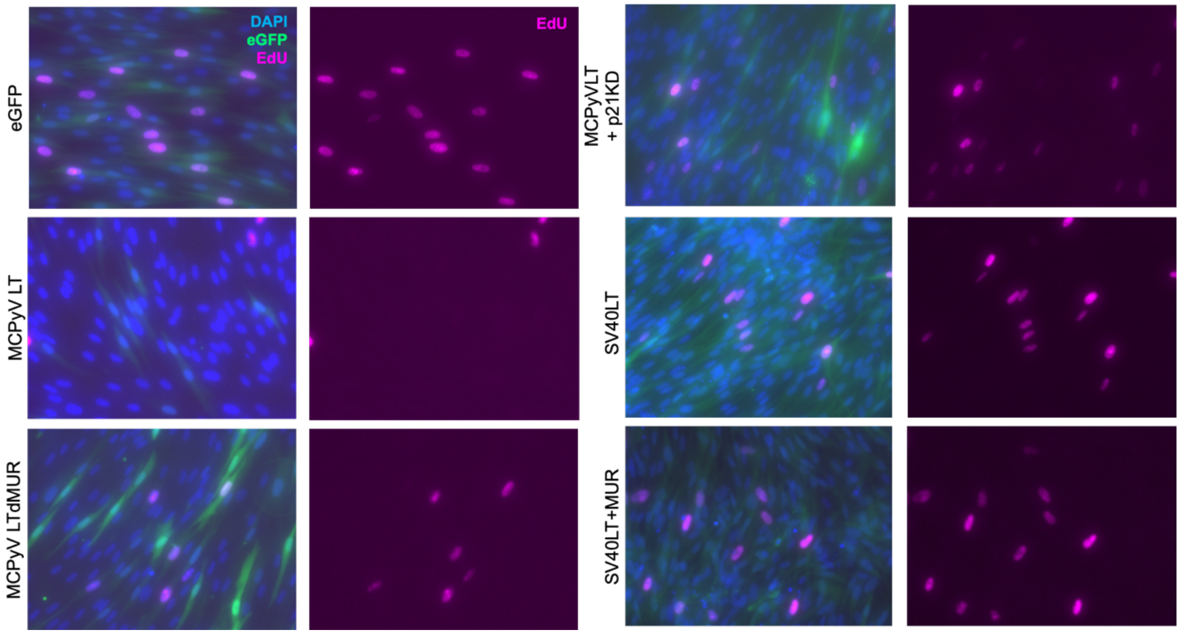


**Fig. S6. MCPyV LT negatively regulates cell growth by inducing cellular senescence.** Immunofluorescent imaging of EdU incorporation is shown. EdU incorporation (pink) was detected, stained with DAPI (blue), and then visualized with a REVOLVE4 fluorescence microscope.

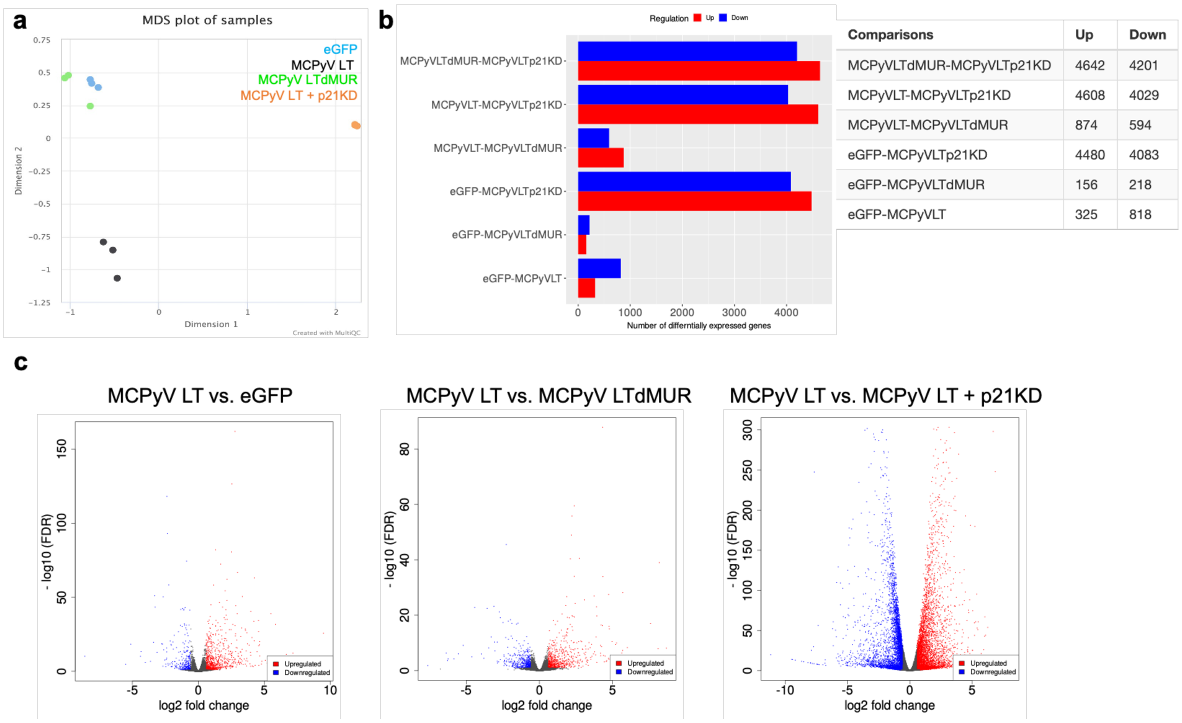


| Comparisons | Up | Down |
|---|---|---|
| MCPyVLTdMUR-MCPyVLTp21KD | 4642 | 4201 |
| MCPyVLT-MCPyVLTp21KD | 4608 | 4029 |
| MCPyVLT-MCPyVLTdMUR | 874 | 594 |
| eGFP-MCPyVLTp21KD | 4480 | 4083 |
| eGFP-MCPyVLTdMUR | 156 | 218 |
| eGFP-MCPyVLT | 325 | 818 |

**Fig. S7: Differential gene expression analysis. (a)** Multidimensional analysis plot illustrating transcriptional profiles between MCPyV LT constructs and replicates. **(b)** Differences in gene expression between samples are shown by histograms displaying the number of upregulated or downregulated genes. **(c)** Volcano plots display upregulated and downregulated differentially expressed genes (DEGs) between indicated samples. False discovery rate (FDR) cutoff was set to 0.1 and $\log_2$ fold change cutoff was set to 1.5.

a

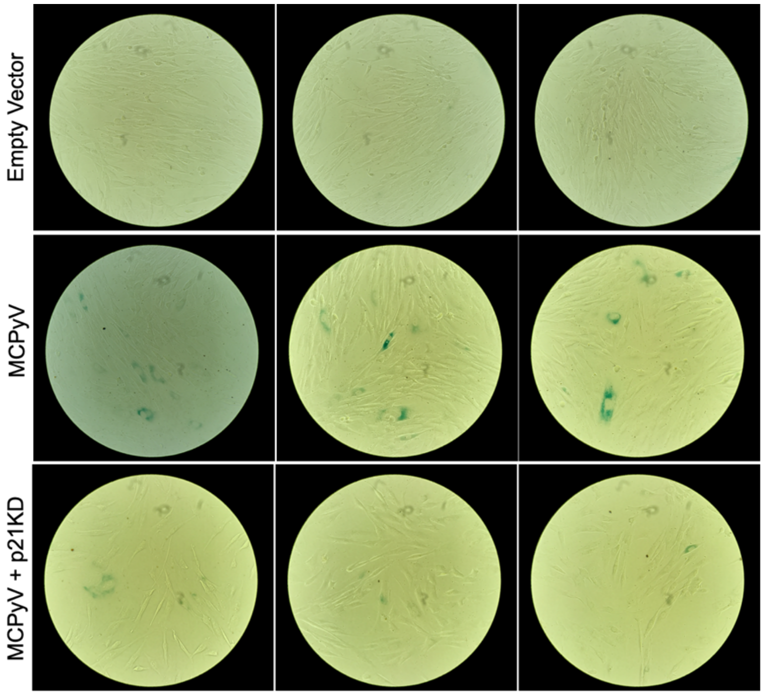


b

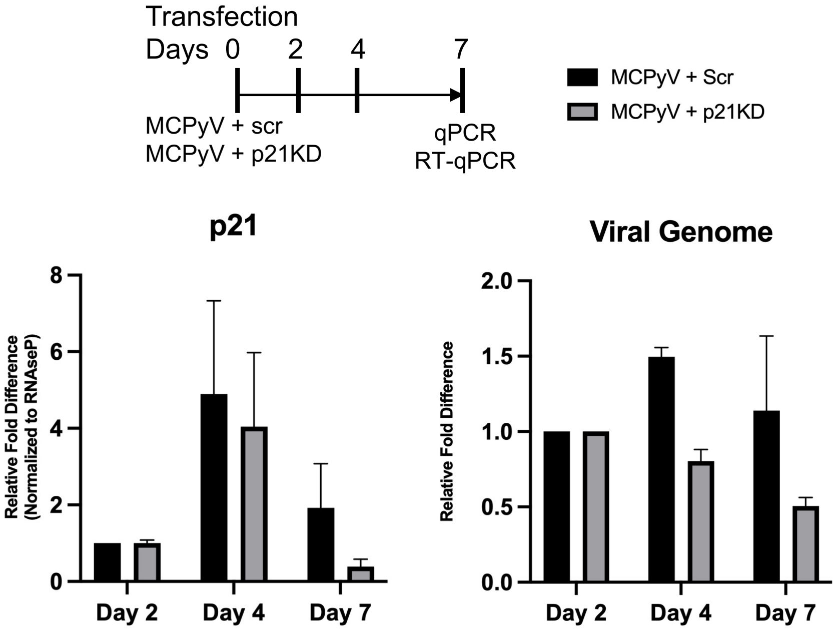


**Fig. S8. MCPyV induces p21-dependent senescence. (a)** SA-β-gal assay. SA-β-gal staining in empty vector control or MCPyV transfected cells with or without p21 knockdown was visualized and photographed under phase contrast and bright field microscopy. At 14 days post transfection, p21 knockdown was performed by lentiviral shRNA transduction for 7 days. **(b)** p21 knockdown reduces MCPyV genome levels. BJ-hTERT cells were cotransfected with MCPyV and mCherry-shScr or mCherry-shp21.1. Cells were harvested at stated days post transfection and were analyzed by RT-qPCR and quantitative PCR (qPCR) to verify knockdown of p21 transcript levels (left) and determine MCPyV genome levels (right) respectively.

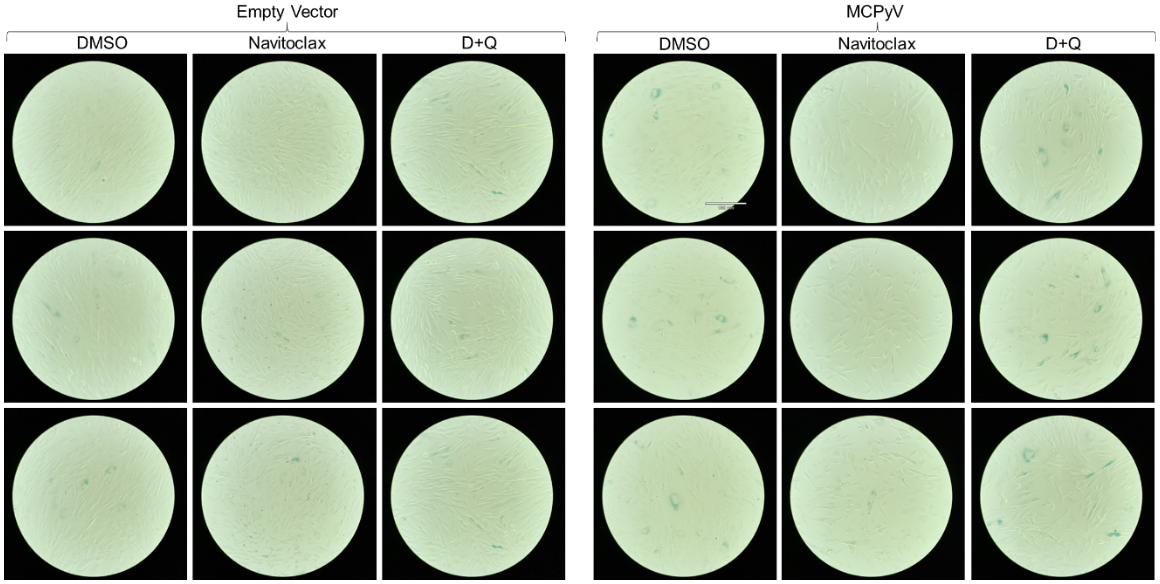


**Fig. S9. Senolytic treatment inhibits MCPyV-induced cellular senescence.** SA-β-gal assay. Empty vector (left) or MCPyV (right) transfected BJ-hTERT cells were grown for 14 days. Cells were then treated with DMSO, navitoclax (250 nM), or dasatinib+quercetin (D+Q, 25 nM dasatinib + 250 nM quercetin) for two days and then subjected to SA-β-gal staining. Senolytics selectively cleared senescent cells induced by MCPyV. Navitoclax treatment substantially reduced MCPyV-induced senescence. Representative images are shown. Scale bar = 180 μm.

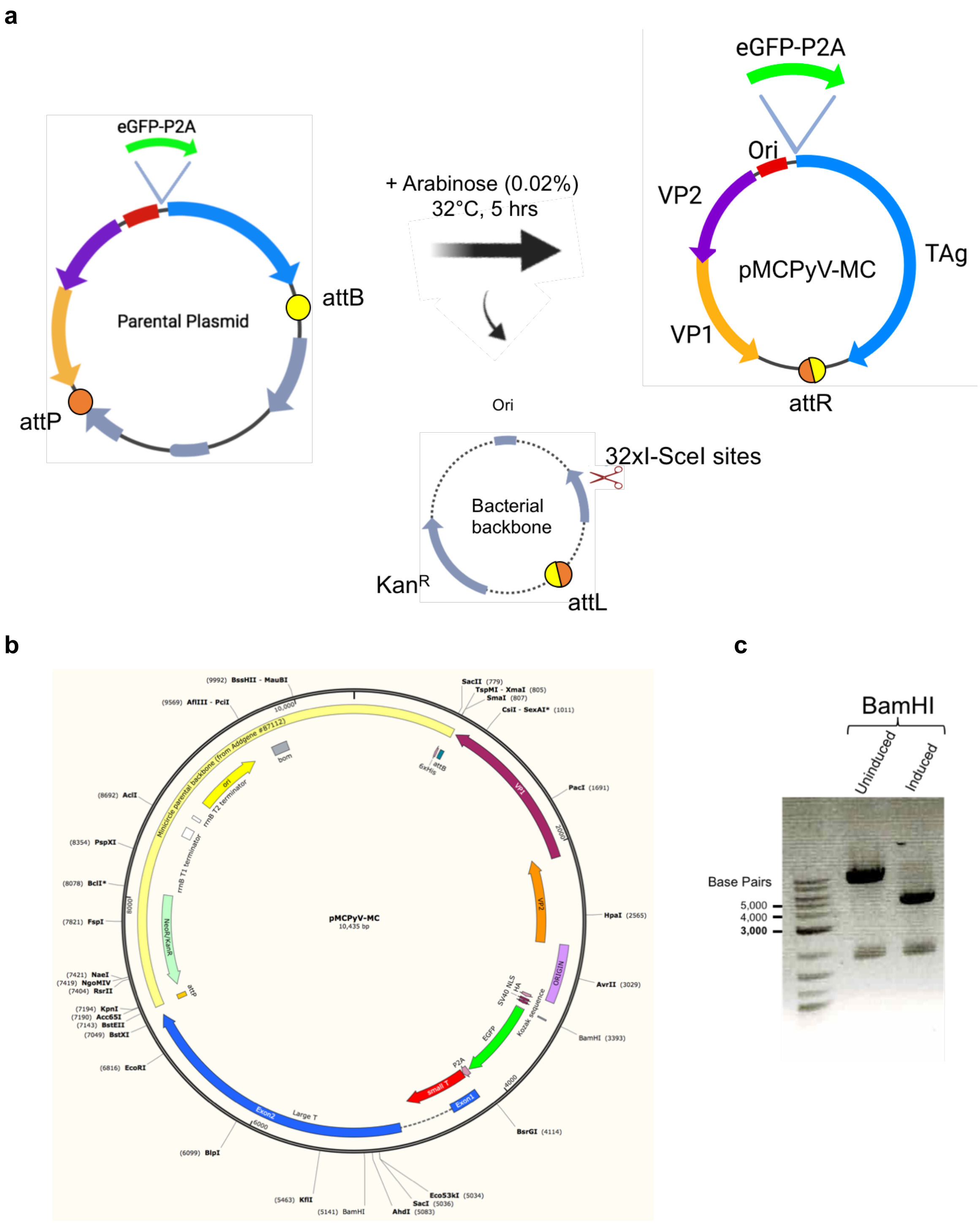


**Fig. S10. MCPyV minicircle genome.**
**(a)** Summarized representation of the strategies used for minicircle purification. **(b)** A construct of MCPyV minicircle genome. **(c)** Electrophoresis of MCPyV minicircle clone. MCPyV genome was isolated before and after induction by adding 0.02% arabinose and digested with BamHI.

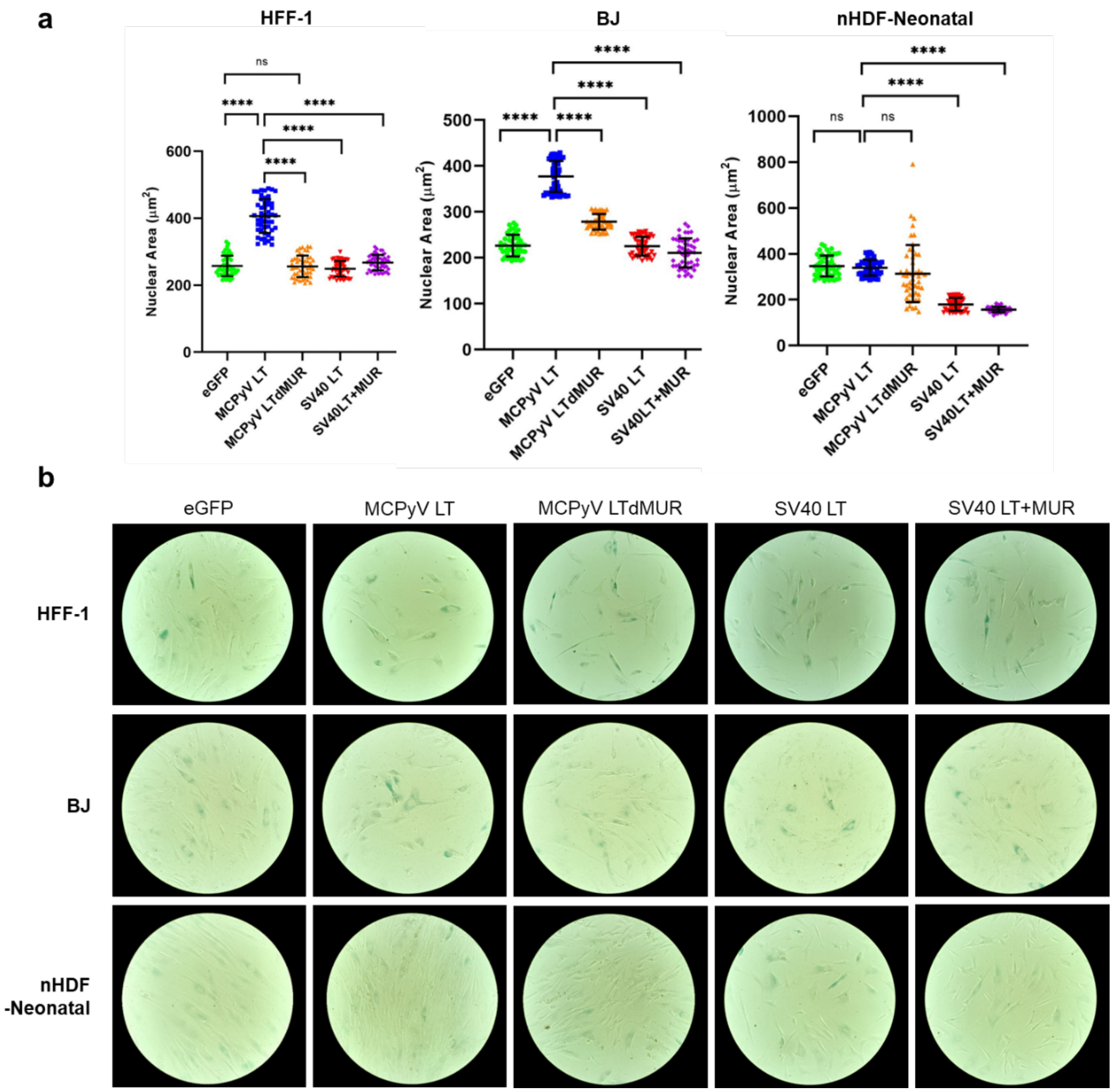


**Fig. S11. MCPyV LT expression exhibits senescence-specific enlarged nucleus in HFF-1 and BJ foreskin fibroblasts but not in primary normal human dermal fibroblast. (a)** Abnormal nuclear size change in MCPyV LT-expressing (GFP-positive) HFF-1 (at passage 13), BJ (at passage 9) and primary normal human dermal fibroblast (nHDF-Neonatal) cells (at passage 5, Lonza). nHDFs are grown using FGM-2 growth medium-2 BulletKit$^{TM}$ (Lonza). Nuclear size was measured 7 days after transduction of each lentiviral construct. Statistical significance was determined using the one-way ANOVA test. Standard error bars represent mean value with standard deviation, n = 50 cells. **(b)** Normal human fibroblast cell lines displayed high senescence-associated β-galactosidase (SA-β-Gal) activity. Thus, BJ-hTERT cell line was used to determine accurate gene changes in MCPyV LT-induced senescence shown in Figure 5.

### Table S1. Primers used in this study.

| Name | Sequence (5' to 3') | Notes |
|---|---|---|
| HFori_AvrII.F | CAACTTGGCTGCCTAGGTG | eGFP-MCPyV-HFwt construct |
| Ori_HA_NLS_GFP11.F | GCATATAGACAAGACCATGTATCCCTATGACGTG | eGFP-MCPyV-HFwt construct |
| Ori_HA_NLS_GFP11.R | CACGTCATAGGGATACATGGTCTTGTCTATATGC | eGFP-MCPyV-HFwt construct |
| P2A_MCVER.F | GAGAACCCTGGACCTGATTTAGTCCTAAATAG | eGFP-MCPyV-HFwt construct |
| P2A_MCVER.R | CTATTTAGGACTAAATCAGGTCCAGGGTTCTC | eGFP-MCPyV-HFwt construct |
| MCV350.1605-1585 | CAGCATGGCTAAGATAATCAG | eGFP-MCPyV-HFwt construct |
| MCVHFmcF-SmaI | CCCCGGGCGCGGGAATGCATGAAATAATTCTCATAATTCTTGTGTTTGGC | pMCPyV-MC construct |
| MCVHFmcR-BstEII | CTCAAAGGTTACCCCAGTTGGGGCGGGCCCTATTCAGTTAAGTAGGCCCCAG | pMCPyV-MC construct |
| Exon1.R | TCCAAAGGGTGTTCAATTCC | pMCPyV-MC construct |
| eGFPqPCR.F | GCAAGGGCGAGGAGCTGTTCAC | pMCPyV-MC construct |
| CDKN1A p21.F | GACACCACTGGAGGGTGACT | RT-qPCR |
| CDKN1A p21.R | CAGGTCCACATGGTCTTCCT | RT-qPCR |
| CSF2 GM-CSF.F | GGCCCCTTGACCATGATG | RT-qPCR |
| CSF2 GM-CSF.R | TCTGGGTTGCACAGGAAGTTT | RT-qPCR |
| ANKRD1.F | AGT AGA GGA ACT GGT CACTGG | RT-qPCR |
| ANKRD1.R | TGG GCT AGA AGT GTCTTC AGA T | RT-qPCR |
| EDN1.F | CAG CAGTCT TAG GCG CTG AG | RT-qPCR |
| EDN1.R | ACTCTT TAT CCA TCA GGG ACG AG | RT-qPCR |
| CXCL1.F | GAAAGCTTGCCTCAATCCTG | RT-qPCR |
| CXCL1.R | CACCAGTGAGCTTCCTCCTC | RT-qPCR |
| CXCL2.F | AACTGCGCTGCCAGTGCT | RT-qPCR |
| CXCL2.R | CCCATTCTTGAGTGTGGCTA | RT-qPCR |
| IL6.F | CCGGGAACGAAAGAGAAGCT | RT-qPCR |
| IL6.R | GCGCTTGTGGAGAAGGAGTT | RT-qPCR |
| IL7.F | CTCCAGTTGCGGTCATCATG | RT-qPCR |
| IL7.R | GAGGAAGTCCAAAGATATACCTAAAAGAA | RT-qPCR |
| IL8.F | CTTTCCACCCCAAATTTATCAAAG | RT-qPCR |
| IL8.R | CAGACAGAGCTCTCTTCCATCAGA | RT-qPCR |
| shP21.1-F | CCGGCGCTCTACATCTTCTGCCTTACTCGAGTAAGGCAGAAGATGTAGAGCGTTTTTG | shRNA (TRCN0000287021) |
| shP21.1-R | AATTCAAAAACGCTCTACATCTTCTGCCTTACTCGAGTAAGGCAGAAGATGTAGAGCG | shRNA (TRCN0000287021) |
| shP21.2-F | CCGGGACAGATTTCTACCACTCCAACTCGAGTTGGAGTGGTAGAAATCTGTCTTTTTG | shRNA (TRCN0000040126) |
| shP21.2-R | AATTCAAAAAGACAGATTTCTACCACTCCAACTCGAGTTGGAGTGGTAGAAATCTGTC | shRNA (TRCN0000040126) |
| GAPDH.F | CCTCCCGCTTCGCTCTCT | RT-qPCR |
| GAPDH.R | CTGGCGACGCAAAAGAAGA | RT-qPCR |
| RNase P.F | GCGGAGGGAAGCTCATCAG | RT-qPCR, MCPyV qPCR |
| RNase P.R | CTGGCCCTAGTCTCAGACCTT | RT-qPCR, MCPyV qPCR |
| MCV350.1605-1585 | CAGCATGGCTAAGATAATCAG | MCPyV qPCR |
| MCV350.1505-1528 | CAGCAAGTTTTACAAGCACTCCACC | MCPyV qPCR |
| eGFPqPCR.F | GCAAGGGCGAGGAGCTGTTCAC | MCPyV qPCR |
| eGFPqPCR.R | GGTGGCATCGCCCTCGCCCTC | MCPyV qPCR |
| HFligRep.F1 | GCTGCTGCAGAGTTCCTCCTATATG | MCPyV qPCR |
| HFligRep.R | GGCTGCAGATACAATCAAACCTAG | MCPyV qPCR |
| pLKOmChe_BHI.F | CAGGGGGATCCATGGTGAGCAAGGGCGAGG | pLKO.1 mCherry-Puro construct |
| pLKO_KpnI_R | GGTCTTAAAGGTACCGATGCATGGGGTC | pLKO.1 mCherry-Puro construct |
| P2Apuro.F | GAACCCTGGACCTAGCGCTATGACCGAGTACAAGCCC | pLKO.1 mCherry-Puro construct |
| mCherryP2A.F | GACGAGCTGTACAAGGGAAGCGGAGCTAC | pLKO.1 mCherry-Puro construct |
| mCherryP2A.R | GTAGCTCCGCTTCCCTTGTACAGCTCGTC | pLKO.1 mCherry-Puro construct |

**Dataset S1 (separate file).** The comprehensive functional enrichment analysis.
